# Glucose deprivation promotes pseudo-hypoxia and de-differentiation in lung adenocarcinoma

**DOI:** 10.1101/2023.01.30.526207

**Authors:** Pasquale Saggese, Aparamita Pandey, Eileen Fung, Abbie Hall, Jane Yanagawa, Erika F. Rodriguez, Tristan R. Grogan, Giorgio Giurato, Giovanni Nassa, Annamaria Salvati, Alessandro Weisz, Steven M. Dubinett, Claudio Scafoglio

## Abstract

Increased utilization of glucose is a hallmark of cancer. Several studies are investigating the efficacy of glucose restriction by glucose transporter blockade or glycolysis inhibition. However, the adaptations of cancer cells to glucose restriction are unknown. Here, we report the discovery that glucose restriction in lung adenocarcinoma (LUAD) induces cancer cell de-differentiation, leading to a more aggressive phenotype. Glucose deprivation causes a reduction in alpha-ketoglutarate (αKG), leading to attenuated activity of αKG-dependent histone demethylases and histone hypermethylation. We further show that this de-differentiated phenotype depends on unbalanced EZH2 activity, causing inhibition of prolyl-hydroxylase PHD3 and increased expression of hypoxia inducible factor 1α (HIF1α), triggering epithelial to mesenchymal transition. Finally, we identified an HIF1α-dependent transcriptional signature with prognostic significance in human LUAD. Our studies further current knowledge of the relationship between glucose metabolism and cell differentiation in cancer, characterizing the epigenetic adaptation of cancer cells to glucose deprivation and identifying novel targets to prevent the development of resistance to therapies targeting glucose metabolism.

## INTRODUCTION

Rewiring of energy metabolism is a hallmark of cancer, and the increased dependency on glucose is a common characteristic of cancer cells (DeBerardinis & Chandel, 2016; Vander Heiden & DeBerardinis, 2017). Malignancies require increased glucose for quick production of energy as well as for shunting carbon atoms toward macromolecule biosynthesis (Martinez & Scafoglio, 2020). Oncogenic signalling induces metabolic rewiring, creating tumor-specific metabolic vulnerabilities. Kras-induced metabolic rewiring is specifically geared toward increased glycolysis (Kerr *et al*, 2016), and glucose uptake via Glut1 and Glut3 is required for Kras-driven oncogenesis (Contat *et al*, 2020). Much interest and research effort has focused on targeting glucose uptake and glycolysis for therapeutic purposes (Goodwin *et al*, 2017; Hsieh *et al*, 2019; Laussel & Leon, 2020; Zhu *et al*, 2021). However, the clinical results with metabolic therapies targeting glucose have been disappointing, likely due to cancer cell plasticity and adaptation mechanisms (Cargill *et al*, 2021). The complex adaptations induced in cancer cells by glucose restriction are not known. Here, we report the discovery that glucose restriction induces de-differentiation and promotes aggression of lung adenocarcinoma (LUAD) cells.

Recent studies have highlighted how metabolic requirements and vulnerabilities evolve during cancer progression (Faubert *et al*, 2020). We previously identified a metabolic vulnerability specific for early-stage LUAD, which relies on sodium-glucose transporter 2 (SGLT2) for growth and progression (Scafoglio *et al*, 2018). Since early lesions of the LUAD spectrum rely on SGLT2 for glucose uptake, inhibition of this transporter with an anti-diabetes drug, canagliflozin, significantly delays the development of LUAD, prolongs survival, and reduces tumor growth in genetically engineered murine models and patient-derived xenografts (Scafoglio *et al.*, 2018). However, the tumors eventually escape the treatment and undergo a de-differentiation process, suggesting that glucose restriction induces a phenotypic switch in LUADs.

Glutamine restriction in the tumor core is known to cause cancer de-differentiation, due to reduced alpha-ketoglutarate (αKG) (Tran *et al*, 2020). αKG is a key modulator of cell differentiation both in normal development and in cancer (Saggese *et al*, 2020). αKG availability has a direct impact on gene expression, since it is required for the activity of the Jumonji C domain (JMJD)-containing histone demethylases and the ten-eleven-translocation (TET) enzymes, involved in DNA de-methylation (Carey *et al*, 2015b). Therefore, αKG depletion leads to histone and DNA hyper-methylation, with repression of differentiation-related genes.

Multiple histone demethylases depend on αKG for their enzymatic activity (Agathocleous & Harris, 2013; Campbell & Wellen, 2018). However, H3K27 tri-methylation plays a major role in regulating glutamine restriction-dependent cell de-differentiation (Pan *et al*, 2016). H3K27 is methylated by the Polycomb Repressor Complex 2 through the activity of histone methyltransferase Enhancer of Zeste Homolog 2 (EZH2), and it is de-methylated by JMJD3 and UTX. Knockout of UTX promotes Kras-driven LUAD in an EZH2-dependent way (Wu *et al*, 2018), while EZH2 over-expression drives LUAD development independently of Kras (Zhang *et al*, 2016), suggesting that this pathway is relevant in LUAD progression.

Here, we show that glucose deprivation in LUAD causes cancer de-differentiation similar to that caused by glutamine restriction. Glucose deprivation reduces αKG availability, limiting αKG-dependent histone demethylase activity and induces histone hypermethylation, driving LUAD to a poorly differentiated state and a highly aggressive phenotype. Surprisingly, we found that glucose restriction-induced de-differentiation and increased aggressiveness requires activation of the hypoxia-inducible factor 1α (HIF1α) signaling. EZH2 is involved in up-regulation of HIF1α by direct repression of the proline hydroxylase PHD3, which initiates HIF1α degradation in normoxia. HIF1α induces Slug activity and epithelial to mesenchymal transition (EMT), leading to a highly aggressive and metastatic phenotype. Targeting the EZH2/HIF1α/Slug axis pharmacologically potentiates the effect of SGLT2 inhibitors in murine LUAD. Finally, we describe a transcriptional signature regulated by the hypoxia pathway whose expression in human LUAD confers a significantly worse prognosis.

## RESULTS

### 1. Glucose restriction induces de-differentiation in LUAD

We previously showed that pharmacological inhibition of SGLT2 in murine LUAD results in reduction of the tumor burden and improvement of survival (Scafoglio *et al.*, 2018). However, mice eventually develop aggressive tumors and die even during the treatment. To find out mechanisms of resistance, we used our previously developed conditional genetically engineered model (GEMM) driven by Kras^G12D^ mutation and p53 deletion (KP mice) (Scafoglio *et al.*, 2018). In this model lung tumors are induced by intranasal administration of adenovirus encoding the Cre recombinase (AdenoCre). Two weeks after tumor induction, we treated the KP mice with empagliflozin, a selective SGLT2 inhibitor, and we collected the lungs at week 8 post-AdenoCre inhalation for histology. Visual inspection of the hematoxylin and eosin (H&E) staining of lung sections showed that the lesions in the SGLT2-treated mice were more poorly differentiated than those in the placebo group. The KP tumors recapitulated a heterogeneous morphology typical of human LUAD, with co-existance of different components: papillary, acinar, solid, and micropapillary (Fig. EV1A). Pathological quantification performed by a board-certified pathologist (Table EV1) showed that empagliflozin changed the differentiation state of the tumors, significanlty reducing the solid component and increasing the micropapillary component of murine tumors (Fig. 1A). The micropapillary component is an independent predictor of poor prognosis in LUAD (Tsutsumida *et al*, 2007). In addition, some of the tumors in the empagliflozin group had a clear cell component which was completely absent from placebo tumors (Fig. EV1A and Table EV1). Clear cell carcinoma is a very rare variety of LUAD with uncertain prognostic significance (Komiya *et al*, 2019). The pathologic assessment suggested that empaglflozin treatment changed the differentiation state of the tumors. To quantify the effect of glucose restriction on tumor differentiation, we performed immunohistochemistry (IHC) on the KP murine lungs with antibodies against FoxA2 and Ttf-1, two major markers of differentiation and regulators of cell state transitions in LUAD (Li *et al*, 2015; Orstad *et al*, 2022). The tumors treated with empagliflozin showed a significant reduction of Ttf-1 (Fig. 1B-C) and FoxA2 (Fig 1D-E). To confirm this phenotypic switch, we repeated empagliflozin treatment, collecting the tumors at week 6 post-tumor induction, the first time point in which mice develop macroscopically dissectible tumors. RNA was extracted from lung tumors for qPCR analysis. We first examined the expression of lung epithelial cell lineage markers in tumors and observed a significant reduction in surfactant protein C (SpC), a marker of type II alveolar cells (ATII), which are the cell of origin of LUAD (Fig. 1F). We also observed reduced expression of markers of Clara cells (Scgb3A2), type I alveolar cells (Pdpn) and ciliated cells (FoxJ1), although the basal levels of these markers were two orders of magnitude lower than that of SpC (Fig. EV1B). In addition, we analyzed the expression of markers of well-differentiated LUAD (FoxA2 and Ttf1), and of poorly differentiated LUAD (HmgA2). We observed a significant reduction of FoxA2 and Ttf1 and, conversely, an increase of HmgA2 expression in tumors treated with empagliflozin compared with placebo (Fig. 1F). As the tumors have a conditional luciferase transgene, we validated luciferase expression (Fig. EV1C), confirming the proper activation of the conditional model in the collected tumor tissue and showing no difference between the two groups.

**Fig. 1.**
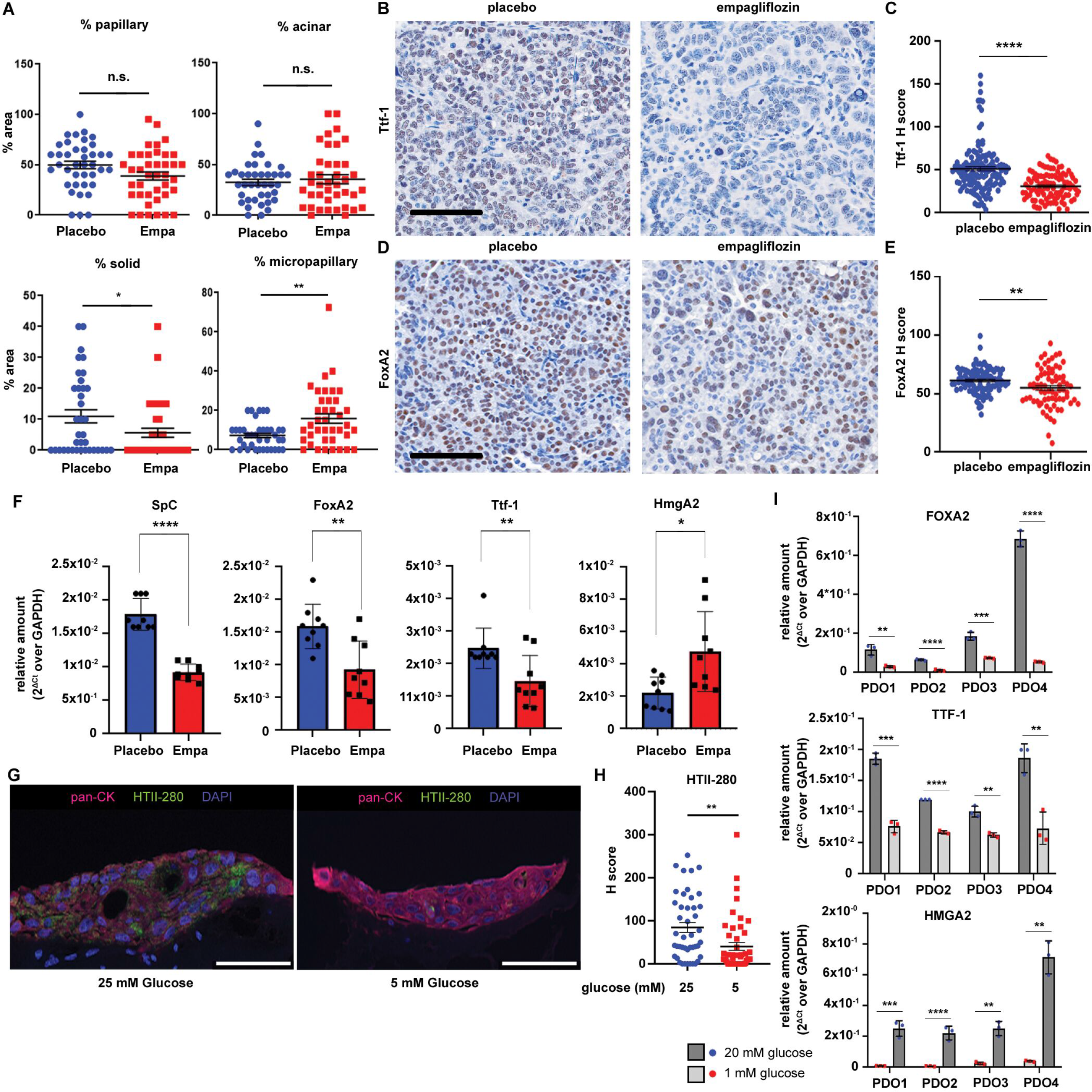
SGLT2 inhibition causes tumor de-differentiation in LUAD. A-F) KP mice carrying LUADs were treated with either placebo or empagliflozin (10 mg/kg/d), starting 2 weeks after tumor induction for 6 weeks. A) Quantification of the percentage of different histological types in hematoxyli and eosin slides from the two treatment groups, performed by a board-certified pathologist. B-E) Representative pictures of IHC stain for Ttf-1 (B) and FoxA2 (D) in the placebo and empagliflozin groups. Quantification of the Ttf-1 (C) and FoxA2 (E) signal, expressed as H score, was measured by QuPath analysis, and significance was estimated by Student’s t-test. Error bars: mean ± SEM. **p < 0.01; ****p < 0.0001. F) KP mice were treated for 4 weeks with either placebo or empagliflozin followed by tumor collection and RNA extraction. RT-PCR was performed to measure the changes in the type II alveolar cell marker SpC and tumor differentiation markers FoxA2, Ttf1, and HmgA2. *p < 0.05; **p < 0.01; ****p < 0.0001. Significance was evaluated by Student’s test. G-I) PDOs were established from fresh surgical specimens of lung adenocarcinoma. All organoids were incubated in high (25 mM), medium (5 mM), or low (1 mM) glucose for three weeks, followed by either immunofluorescence or RT-PCR. G-H) Immunofluorescence with the indicated antibodies was performed in one PDO incubated in high or medium glucose. Representative images (G) and quantification of the signal of three pooled independent experiments (H) are reported. Significance was measured with Student’s T-test. **p < 0.01. I) RNA was extracted from 4 PDOs incubated in high or low glucose for RT-PCR with primers specific for FOXA2, TTF1, and HMGA2. Significance was measured by Student’s T-test. *p < 0.05; **p < 0.01; ***p < 0.001; ****p < 0.0001.

To confirm the effect of glucose deprivation on cell differentiation in a more clinically relevant model, we established patient-derived organoids (PDOs) from a surgical sample of early-stage lung adenocarcinoma. We cultured the newly established PDOs in either high (25 mM) or medium (5 mM) glucose for three weeks, followed by immunofluorescence. The immunofluorescence experiment showed that the expression of HTII-280, a marker of well-differentiated LUAD (Gonzalez *et al*, 2010), was significantly reduced by incubation in 5 mM compared with 20 mM glucose (Fig. 1G-H). Five mM is a lower concentration than the typical cell culture conditions (25 mM); however, this concentration is close to normal blood glucose level. Therefore, we aimed at confirming this result with lower glucose concentrations. We incubated 4 different freshly established PDOs in high (20 mM) or low (1 mM) glucose for three weeks, with addition of mannitol as osmolarity control in the low-glucose samples, and extracted RNA for RT-PCR. This experiment showed that markers of well-differentiated LUAD, FOXA2 and TTF1, were significantly down-regulated in all four PDOs, whereas poor differentiation marker HMGA2 was up-regulated in three out of four PDOs by incubation in low glucose (Fig. 1I).

Taken together, these results suggest that glucose restriction, induced by SGLT2 inhibition or by incubation in low glucose culture, leads the tumors towards a poorly differentiated phenotype.

### 2. Glucose restriction leads de-differentiation in LUAD cell lines and drives a more aggressive phenotype

To investigate more in detail the effects of glucose deprivation on cell differentiation, we performed *in vitro* experiments in two human cell lines, A549 and NCI-H358, which have been described as moderately differentiated (Lieber *et al*, 1976; Nardone & Andrews, 1979; Sumi *et al*, 2018) and express high levels of FoxA2, as well as a murine cell line, 2953A, which was established in our lab from a KP lung tumor. We decided to study the effects of glucose deprivation by incubation of cell lines in low glucose.

To investigate the relationship between glucose metabolism and cell differentiation, we performed liquid chromatography-mass spectrometry (LC-MS) analysis to detect changes in metabolites caused by incubation of A549 and NCI-H358 cells overnight in low glucose. We focused specifically on metabolites involved in glycolysis and the TCA cycle. As expected, low glucose significantly reduced the levels of glycolytic and TCA cycle metabolites in both A549 and NCI-H358 cells, except for Acetyl-CoA, which showed no significant changes. Representative metabolites of the two pathways are presented in Figure 2A (A549 cells) and Fig. EV1D (NCI-H358 cells). A list of all the metabolites is reported in Table EV2.

**Fig. 2.**
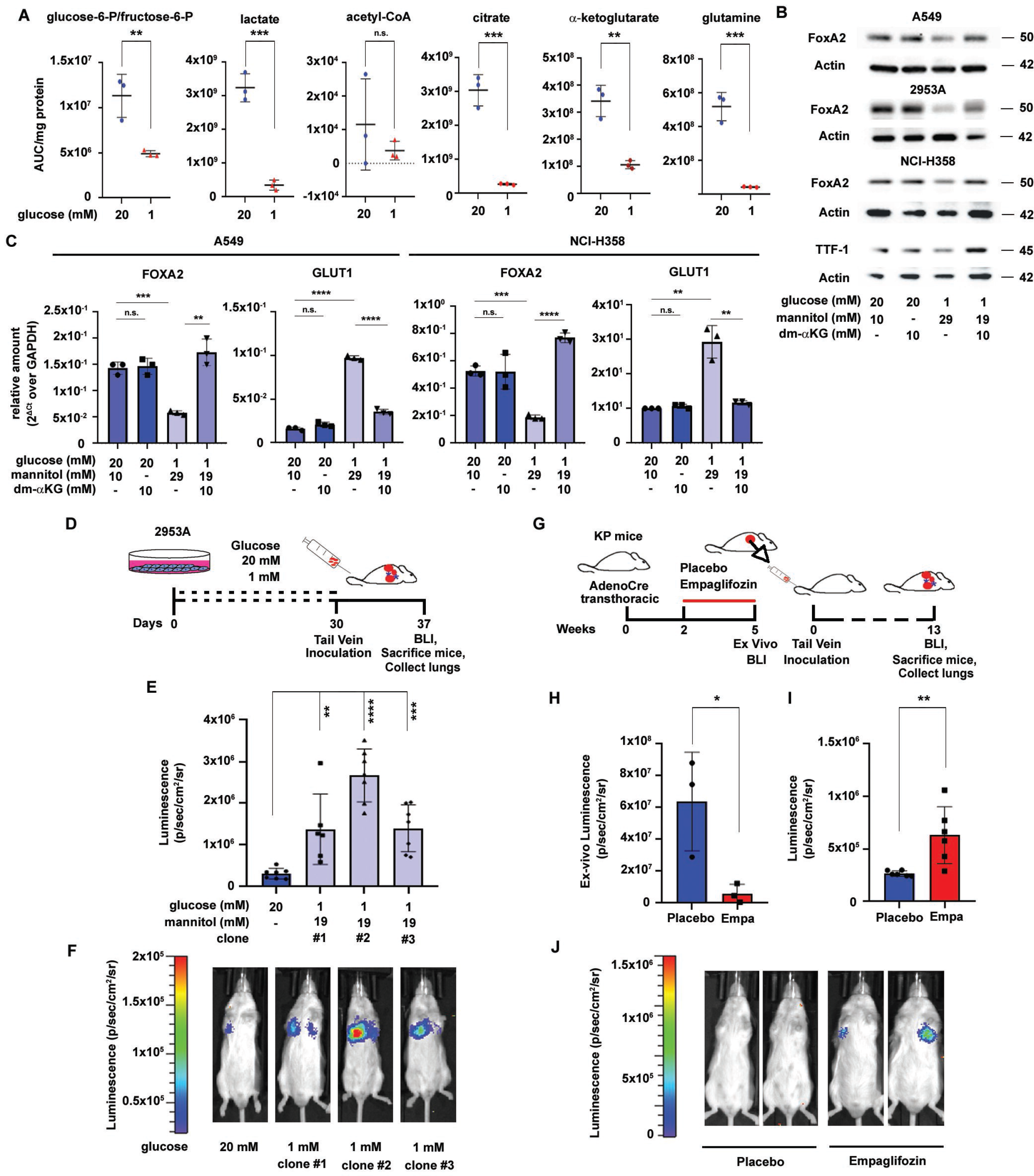
Glucose restriction causes LUAD de-differentiation, due to low αKG and histone hypermethylation, and increases cell aggressiveness. A) A549 cells were incubated in high (20 mM) or low (1 mM) glucose for 24 hours and mass spectrometry analysis was performed. The results are expressed as metabolite amounts (area under the curve – AUC) normalized by total protein amounts. The statistical significance of the observed changes was evaluated by one-way ANOVA: **p < 0.01, ***p < 0.001. B-C) A549, NCI-H358, and 2953A cells were cultured for 5 days in medium containing high or low glucose with or without αKG, as indicated. B) Western blot analysis of FoxA2 and TTF1 (only for NCI-H358, because the other cell lines did not express this marker). β-actin was used as loading control. C) RT-PCR analysis of expression of FOXA2 and GLUT1 in NCI-H358 cells. GAPDH was used as housekeeping gene. **p < 0.01; ***p < 0.001; ****p<0.0001. D-F) Murine LUAD cells 2953A were incubated in either high (20 mM) or low (1 mM) glucose for at least one month (D). Three clones from cells incubated in low glucose were picked and cultured separately. Both high-glucose and low-glucose cells were inoculated in syngeneic mice by tail vein injection to measure the development of lung metastases. The tumor burden was measured by bioluminescence imaging (BLI). Both quantification of the signal (E) and representative pictures of single mice (F) are reported. G-L) KP mice carrying LUADs were treated with placebo and empaglifozin (10 mg/kg/d), starting 2 weeks after tumor induction by transthoracic injection of AdenoCre. At week 5 post-tumor induction, mice (n = 3) were sacrificed for tumor isolation, single cell dissociation, sorting of epithelial cells, and i.v. injection in syngeneic mice (G). Tumor burden was estimated by BLI for 13 weeks after re-injection. H) Quantification of *ex vivo* BLI at week 5 after the end of the treatment trial. I) Quantification of the *in vivo* BLI at week 13 after re-injection in syngeneic mice. L) Representative images of BLI at week 13. Significance was evaluated by Student’s t-test. *p<0.05; **p<0.01; ***p < 0.001; ****p < 0.0001.

To test the effect of glucose depletion on LUAD differentiation markers, we performed a time-course experiment by growing the A549 and NCI-H358 cells in low glucose (1 mM) for up to 9 days. We observed that glucose restriction down-regulated the expression of well-differentiated marker FoxA2 (Fig. EV1E) and up-regulated the expression of poor differentiation marker HmgA2 (Fig. EV1F) after 5 days of treatment, confirming that glucose restriction induces LUAD de-differentiation.

Alpha-ketoglutarate (αKG) plays a fundamental role on cell differentiation because it is a co-factor of histone and DNA demethylases (Carey *et al*, 2015a). Depletion of glutamine causes cancer de-differentiation due to reduced αKG (Pan *et al.*, 2016) and consequent inhibition of αKG-dependent demethylases, and DNA/histone hyper-methylation. To test if the reduction in FoxA2 in low glucose was due to αKG depletion, we incubated A549, NCI-H358, and 2953A cells in low glucose for 5 days, with or without addition of the cell membrane-permeable precursor dimethyl-αKG (dm-αKG, 10 mM), which gets converted to αKG in the cytosol (Doucette *et al*, 2011). Furthermore, we measured expression of another well-differentiated LUAD marker, TTF1, in NCI-H358 cells, whereas A549 and 2953A cells did not express this marker. We used mannitol as an osmolarity control in low-glucose culture conditions. As expected, incubation in low glucose caused a consistent reduction in FOXA2 and TTF1, rescued by dm-αKG (Fig. 2B). Analysis of mRNA levels by qPCR showed a significant inhibition of TTF-1 in NCI-H358 cells (Fig. EV1G), and of FoxA2 in A549, NCI-H358 (Fig. 2C), and 2953A cells (Fig. EV1H), rescued by dm-αKG. In addition, low glucose increased GLUT1 expression, and dm-αKG reduced its levels (Fig 2C and EV1H). We have previously shown that GLUT1 is expressed in poorly differentiated LUADs (Scafoglio *et al.*, 2018). GLUT1 can be up-regulated by glucose restriction in an αKG- and NF-kB-dependent way (Wang *et al*, 2019). GLUT1 up-regulation can provide an alternative mechanism of glucose supply in cancer cells, conferring resistance to SGLT2 inhibitors.

Since poorly differentiated tumors are typically more aggressive than well-differentiated lesions, we investigated whether glucose deprivation caused a shift toward a more aggressive phenotype in LUAD. To this aim, we incubated the 2953A cell line, which has transgenic luciferase expression, for one month in either high (20 mM) or low (1 mM) glucose. We expanded three clones of low-glucose growing cells, followed by tail vein inoculation in syngeneic mice (Fig. 2D). One week post-injection, we detected that the high-glucose cells developed very small tumor burden as estimated by bioluminescence imaging (BLI), whereas the low-glucose cells showed significantly higher BLI signal (Fig 2E-F). To test if also *in vivo* SGLT2 inhibition can increase the aggressiveness of cancer cells, we treated KP mice with empagliflozin from week 2 to week 5 after tumor induction by transthoracic injection of AdenoCre, followed by tumor collection, dissociation into single cells, sorting of epithelial cells, and re-inoculation in syngeneic mice by tail vein injection (Fig. 2G). Ex vivo BLI showed that treatment with empagliflozin significantly reduced the tumor burden at week 5 (Fig. 2H); however, when the same number of placebo and empagliflozin-treated cells were injected in syngeneic mice, the empagliflozin-treated cells formed larger tumors than the placebo group (Fig. 2I-J). These data showed that glucose starvation, while slowing tumor growth for reduced proliferation (Scafoglio *et al.*, 2018), eventually induces a more aggressive phenotype in LUAD cells.

### 3. The histone H3K27 is required for cancer de-differentiation in response of glucose deprivation

Previous studies have shown that glutamine deprivation causes cancer de-differentiation due to unbalanced histone methylation (Pan *et al.*, 2016; Tian *et al*, 2013). To assess the effect of glucose deprivation on histone modification, we incubated A549, NCI-H358 and 2953A cells in high (20 mM), medium (5 mM), and low (1 mM) glucose for 5 days, followed by histone extraction and western blot for histone marks. We observed increased tri-methylation of H3K4, H3K9 and H3K27 as the glucose concentration was reduced in all three cell lines. Conversely, H3K27 acetylation was progressively reduced by low glucose (Fig EV2A). Interestingly, the administration of dm-αKG was able to reverse the H3K4, H3K9, and H3K27 hyper-methylation and the H3K27 hypo-acetylation (Fig. 3A). Since H3K27me3 has a major role in regulating glutamine restriction-dependent cancer de-differentiation (Pan *et al.*, 2016), we focused on this histone mark. To confirm that glucose restriction caused H3K27 hyper-methylation in LUAD cells *in vivo*, we induced tumors by transthoracic injection of AdenoCre and treated KP mice with empagliflozin for 1 week, followed by collection of tumors and ELISA assay for H3K27me3. We observed that SGLT2 inhibition caused a significant increase of tri-methylation on H3K27 compared with the placebo group (Fig. 3B). Since EZH2 is the only known methyltransferase targeting H3K27, these results suggest that glucose restriction causes αKG reduction and deficient activity of αKG-dependent histone demethylases, resulting in an unbalanced activity of methyltransferase EZH2 and hyper-methylation on H3K27.

**Fig. 3.**
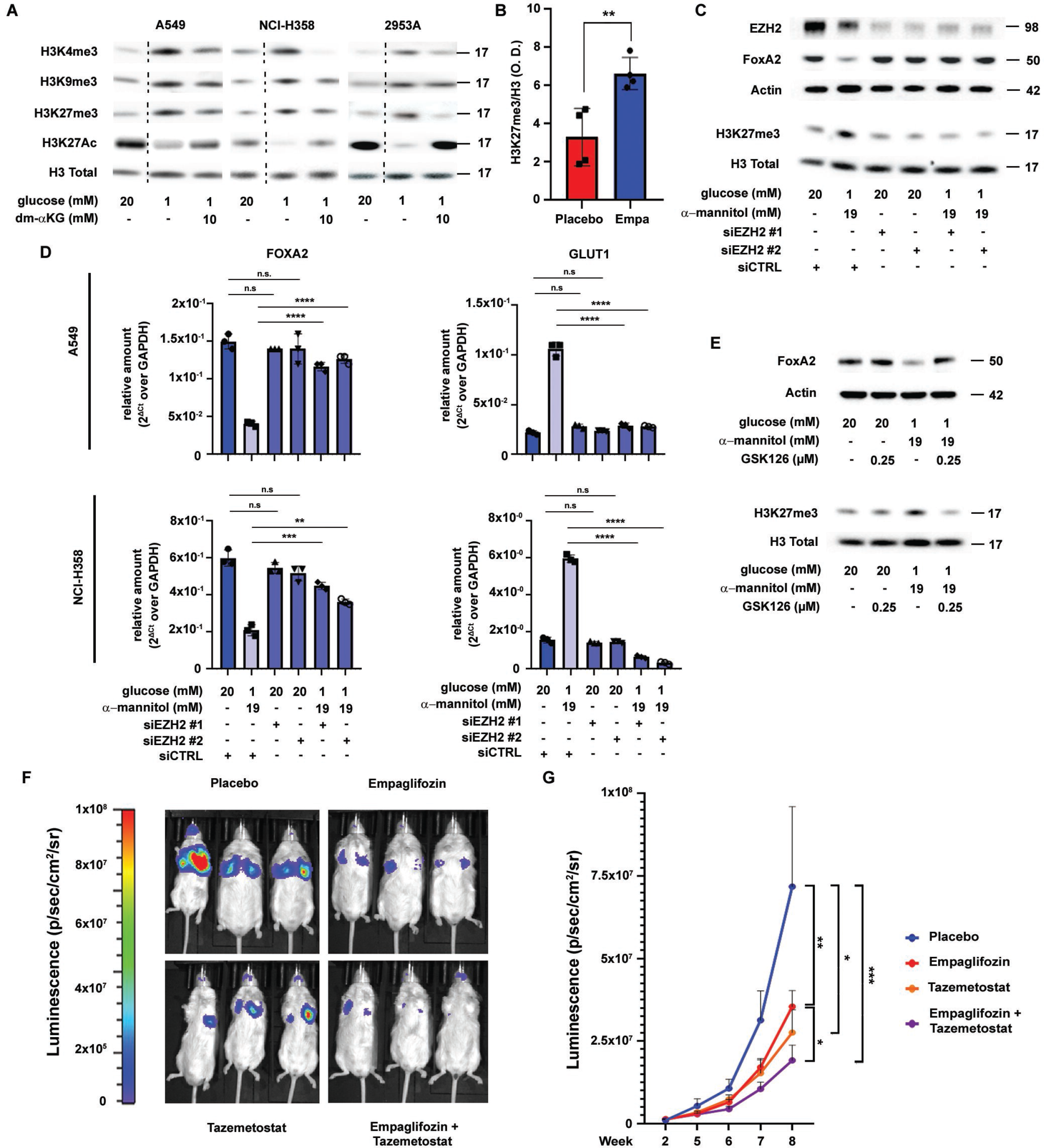
EZH2 is required for cell de-differentiation induced by glucose restriction. A) A549, NCI-H358, and 2953A cells were cultured for 5 days in medium containing high or low glucose with or without dm-αKG, as indicated. Histone marks were analyzed on histone extracts by western blot, as indicated. Total histone 3 (H3) was used as loading control. B) KP mice were treated with empagliflozin (10 mg/kg/d) for 4 weeks starting at week 2 post-tumor induction. The tumors were collected and used for ELISA assay for quantification of H3K27me3. The results are expressed as ratio of H3K27me3 over total histone 3. C-E) A549 cells were cultured for 5 days in RPMI containing either high (20 mM) or low glucose (1 mM), with or without transfection of siRNAs targeting EZH2 (C-D) or treatment with EZH2 inhibitor GSK126 (E). C) Western blot analysis of FOXA2 expression, as well as confirmation of siRNAs efficacy by western blot for EZH2 on whole cell extracts and H3K27me3 on histone extracts. Actin is the loading control for the whole cell extracts and H3 for the histone extracts. D) RT-PCR for FOXA2 and GLUT1. Significance was evaluated by Student’s t-test. **p < 0.01; ***p < 0.001; ****p < 0.0001; n.s.: not significant. E) Western blot for FOXA2 and H3K27me3 in cells treated with GSK126. F-G) KP mice were enrolled in a treatment trial with four groups: 1) placebo; 2) empagliflozin (10 mg/kg/d by oral gavage); 3) tazemetostat (125 mg/kg/bid by oral gavage); 4) empagliflozin + tazemetostat. Treatment was started 2 weeks after tumor induction by intranasal inhalation, and continued until week 8 post-induction. Tumor burden was estimated by bioluminescence imaging: representative images (F) and quantification in all the groups with pooled two biological replicated (G) are reported. Significance was measured using generalized estimating equation models (LIANG & ZEGER, 1986) with terms for time, group, and time by group interaction. *p < 0.05; **p < 0.01; ***p < 0.001

To confirm the role of EZH2 in glucose restriction-induced de-differentiation, we used siRNAs. We first tested the efficiency of two different siRNAs targeting EZH2 separately by RT-PCR (Fig. EV2B). We incubated A549 and NCI-H358 cell lines in either high or low glucose and transfected the cells with either siRNAs targeting EZH2 or control siRNA, followed by protein extraction and Western blot analysis. As expected, incubation in low glucose reduced the expression of FoxA2 and H3K27me3 mark in both cell lines, and EZH2 knockdown rescued FoxA2 expression and H3K27 methylation in low glucose, whereas EZH2 knockdown did not have any effect in the cells incubated in high glucose (A549: Fig. 3C; NCI-H358: Fig. EV2C). RT-PCR analysis confirmed that low glucose incubation reduced expression of FoxA2 and increased the expression of Glut1 in both cell lines, and both were rescued by EZH2 knockdown, whereas the siRNAs targeting EZH2 had no effect on the basal levels in high glucose (Fig. 3D). Finally, we incubated the two cell lines in high or low glucose, with additional inhibition of EZH2 via GSK126, a selective EZH2 inhibitor (Zeng *et al*, 2017). We tested different concentrations (Fig. EV2D) and we observed that a concentration of 0.25 μM inhibited EZH2 activity and rescued the expression of FoxA2 and tri-methylation of H3K27 in both cell lines (A549: Fig 3E; NCI-H358: Fig. EV2E). Overall, these data confirm that EZH2 plays an important role in glucose restriction-induced cancer cell de-differentiation.

Our data suggest that glucose deprivation causes de-differentiation due to insufficient activity of αKG-dependent histone de-methylases. To confirm this hypothesis, we performed RNA interference experiments with siRNAs targeting the two histone de-methylases active on H3K27: UTX and JMJD3. Western blot analysis showed that knockdown of each of these de-methylases in high glucose culture can mimic the effect of low glucose, causing reduction in FoxA2 levels in both A549 and NCI-H358 cells (Fig. EV2F), confirming the importance of H3K27me3 on the maintenance of cell differentiation, and suggesting that both de-methylases are required for cell differentiation in LUAD.

Therefore, we sought to test the hypothesis that EZH2 inhibition, by preventing glucose restriction-induced de-differentiation, would potentiate the effect of SGLT2 inhibition in LUAD. We treated KP mice with SGLT2 inhibitor empagliflozin, EZH2 inhibitor tazemetostat, or both for 6 weeks starting at week 2 post-tumor induction, following the tumor burden by bioluminescence imaging. This trial showed that both empagliflozin and tazemetostat significantly reduced the tumor burden compared with the placebo group, and the combination treatment significantly reduced the tumor burden compared with empagliflozin single agent (Fig. 3F-G). These data confirm that EZH2 inhibition has an additive effect to SGLT2 inhibitor therapy.

### 4. Glucose restriction changes gene expression profiles in LUAD cells, inducing cell proliferation block and dysregulation of cellular differentiation

Since we observed an important role of glucose in maintaining cell differentiation in LUAD, to find the pathways regulated by glucose deprivation, we analyzed the effect of glucose deprivation on gene expression profiles in A549 and NCI-H358 cells. We incubated both cell lines in high glucose, low glucose, or low glucose with supplementation of dm-αKG for five days, followed by RNA-seq analysis. Incubation in low glucose caused up-regulation of 2,394 genes in A549 cells and 1,335 genes in NCI-H358 cells, and down-regulation of 2,024 genes in A549 cells and 942 genes in NCI-H358 cells (Table EV3). Of these, 518 were consistently up-regulated and 439 down-regulated in both cell lines (Fig. EV3A). Addition of dm-αKG to the low glucose culture rescued the expression of the majority of glucose-dependent genes in A549: 1,634 up- and 1,431 down-regulated genes were rescued by dm-αKG (Fig. EV3B). Fewer genes were rescued in H358 cells: 510 up-regulated genes and 370 down-regulated genes (Fig. EV3C). We sought to find genes that were commonly regulated by low glucose and rescued by dm-αKG in the two cell lines, without excluding from the analysis genes that were barely below the threshold of significance. Therefore, we decided to continue our analysis on genes that were significantly regulated in at least 3 of the four conditions (low glucose vs high glucose, low glucose + dm-αKG vs low glucose, in both cell lines) (Table EV3, where the Class number indicates the number of conditions in which the gene is significantly regulated). A minority of genes (75 in total) were regulated discordantly by low glucose in the two cell lines, and we excluded these from our further analyses. We identified 524 genes up-regulated and 445 genes down-regulated by low glucose consistently in both cell lines and rescued by dm-αKG. We focused our analysis on these glucose-regulated, αKG-dependent 969 genes.

Fig. 4A shows volcano plots of the glucose-regulated, αKG-dependent genes in A549 and NCI-H358 cells. Visual inspection of the plots showed that the most down-regulated genes in both cell lines were involved in cell cycle and mitosis (MCM genes, E2F8, ESCO2, SPC25, RRM2, KIF20A, CDC20, and several histone subunits). Conversely, the most up-regulated genes included genes involved in neuronal differentiation (BEX2, PCDH1, AKNA, UNC5B, CUX1, FLRT1, PSAP, APP, SPX, PHRP), hematopoietic differentiation (SLFN5, LCN2), and hypoxia-regulated genes (NUPR1, UNC5B, LAMP3, RRAGD, GDF15, DNAJC2, DDIT3), suggesting that low glucose stopped cell proliferation and mitosis, along with dysregulation of cell differentiation and activation of hypoxia signaling. Consistently, Ingenuity Pathway Analysis (IPA) showed inhibition of functions related to cell proliferation (Cell Cycle Control of Chromosomal replication, Kinetochore Metaphase signaling Pathway, Cyclins and Cell Cycle Regulation, Cell Cycle Regulation by BTG Family Proteins), DNA repair (Nucleotide Excision Repair), as well as epithelial cell identity (Epithelial Adherens Junction Signaling). Conversely, IPA showed activation of stress pathways (Senescence Pathway, Endoplasmic Reticulum Stress Pathway, NRF2-mediated Oxidative Stress Response, Autophagy, Unfolded Protein Response), pathways related to neuronal differentiation (Neuregulin Signaling, Synaptic Long-Term Depression) and hematopoietic differentiation (Fcγ Receptor-mediated Phagocytosis in Macrophages and Monocytes, PI3K Signaling in B Lymphocytes), and hypoxia (HIF1α Signaling) (Fig. 4B and Table EV4). Fig. 4C shows representative down-regulated genes associated with epithelial cell identity and cell cycle, and up-regulated genes associated with stress and hypoxia signaling. Consistently with the IPA results, Gene Set Enrichment Analysis showed down-regulation of cell proliferation pathways (E2F Targets, G2/M checkpoint, and others), and up-regulation of stress pathways (TNFα signaling via NFKB, Unfolded Protein Response, Inflammatory response), as well as cell stemness (Hedgehog Signaling), oncogenic (KRAS Signaling), and hypoxia pathways (Fig. 4D and Table EV5). We confirmed by RT-PCR the low glucose-dependent down-regulation of some genes associated with epithelial cell identity (TUBB4A, TUBB4B) and up-regulation of hypoxia-dependent genes (VEGFB, HK2) (Fig. 4E). Overall, RNA-seq analysis suggested that glucose restriction caused a proliferation block and activation of stress signals in LUAD cell lines, associated with dysregulation of cell differentiation, with down-regulation of epithelial markers and up-regulation of genes associated with neuronal and hematopoietic lineages, as well as the hypoxia pathway. These observations were very interesting because quiescence and increased response to the stress and hypoxia can be associated to resistance to therapy (De Angelis *et al*, 2019).

**Fig. 4.**
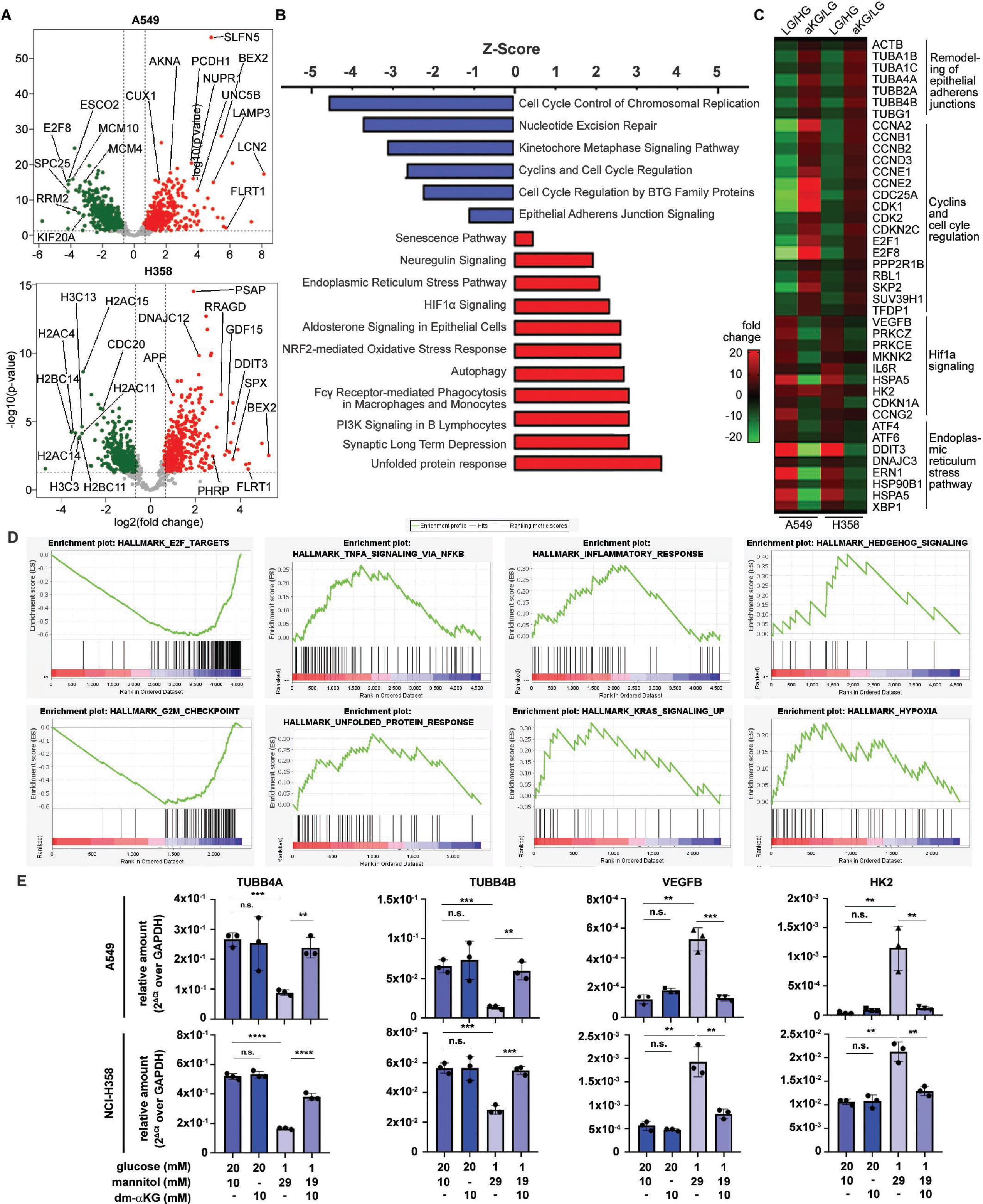
Glucose restriction affects gene expression patterns in LUAD cells. A549 and NCI-358 cells were incubated in high (HG), low (LG) glucose, or low glucose plus αKG (aKG) for 5 days, followed by RNA extraction of RNA-seq analysis. We focused our analysis on the genes that were commonly up* or down-regulated in both cell lines and that were rescued by αKG. A) Volcano plots of genes up- and down-regulated in both cell lines. B) Representative pathways identified by IPA in the genes up* or down-regulated by glucose in both cell lines. C) Heat map of selected genes up* or down-regulated by low glucose and rescued by αKG in both cell lines. D) Representative GSEA plots of pathways enriched in low glucose vs high glucose. E) RT-PCR analysis was performed on selected genes identified from gene expression analysis in A549 and NCI-H358 cell lines incubated in high (20 mM) and low (1 mM) glucose with or without the supplement of αKG. α-mannitol was used as osmotic control.

### 5. Glucose restriction induces shifts in chromatin marks associated with epithelial to mesenchymal transition

Since we observed an important role of EZH2 in glucose restriction-induced de-differentiation, we decided to investigate the global distribution of the H3K27me3 mark in A549 cells incubated for 5 days in low (1 mM) vs high (20 mM) glucose by chromatin immunoprecipitation-sequencing (ChIP-seq). As expected, incubation in low glucose caused a massive repositioning of H3K27me3 in the genome: 12,662 sites present in high glucose were not present in low glucose; conversely, 9,773 sites were present in low glucose but not in high glucose (Fig. 5A). A minority of the sites (5,538) were present both in high and low glucose. Fig. 5B shows a heat map with changes in H3K27me3 genomic locations in high vs low glucose. A panoramic analysis of the localization of these H3K27me3 sites in the genome showed that most of these sites were intronic and intergenic (Fig. 5C). Table EV6 reports a more detailed description of the intergenic sequences with enriched H3K27me3 in high and low glucose, including repetitive sequences (LINE and SINE), long terminal repeats (LTR), and other non-specified intergenic regions. H3K27 methylation in large intergenic regions has been associated with X chromosome inactivation (Zylicz *et al*, 2019) and with silencer elements (Cai *et al*, 2021). We next wanted to analyze the changes in H3K27me3 on the genes that were regulated by low glucose in A549 cells in the RNA-seq experiment. As expected, the H3K27me3 mark was significantly increased in the genes down-regulated by low glucose (Fig. 5D), whereas there was no change in the genes that were up-regulated by low glucose (Fig. 5E). These results confirmed that changes in EZH2 activity are prevalent in the cellular response to glucose deprivation, and are involved in silencing of genes associated with cell proliferation and epithelial differentiation.

**Fig. 5.**
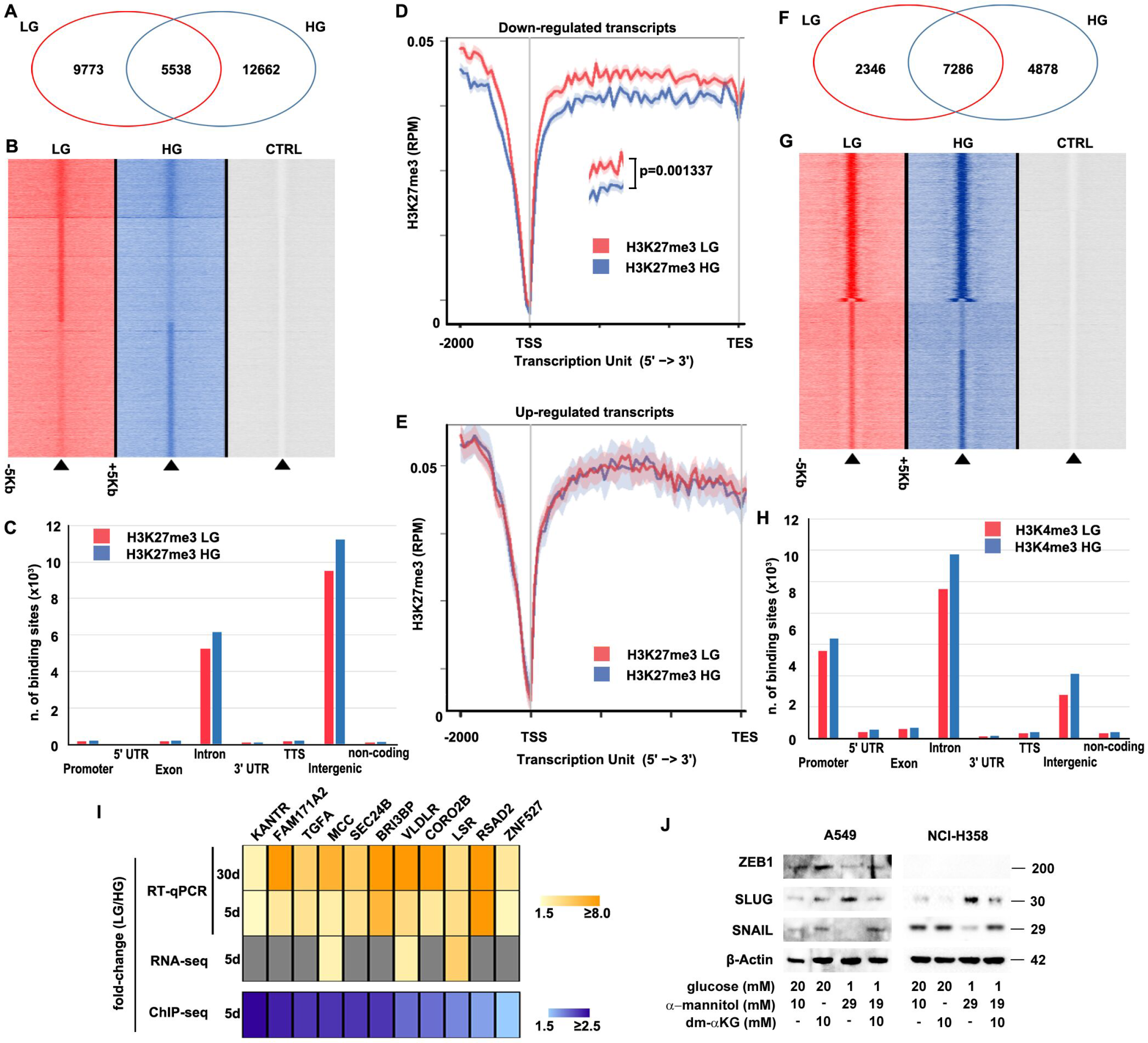
Glucose restriction causes repositioning of H3K27me3 on repressed genes and enrichment of H3K4me3 on Slug target genes. A549 cells were incubated in either high (HG) or low (LG) glucose for 5 days, followed by immunoprecipitation with specific antibodies targeting H3K27me3 and H3K4me3. A-B) Global distribution of the H3K27me3 mark in high vs low glucose, expressed as Venn diagram (A) and heatmap (B). C) Functional classification of the H3K27me3-enriched sites in high and low glucose. D-E) Changes in H3K27 trimethylation in genes that were down-regulated (D) or up-regulated (E) by low glucose in the RNA-seq experiment. F-G) Global distribution of the H3K4me3 mark in high vs low glucose, expressed as Venn diagram (F) and heatmap (G). H) Functional classification of the H3K4me3-enriched sites in high and low glucose. I) Comparison of ChIP-seq, RNA-seq, and RT-qPCR analysis at 5 days and at 30 days on a subset of 11 genes which showed significant increase in H3K4me3 mark in low vs high glucose. J) Western Blot analysis of EMT transcription factors SNAIL, SLUG and ZEB1 in A549 and NCI-H358 cells incubated for 5 days in medium containing different concentrations of glucose with or without αKG, as indicated.

Since the RNA-seq analysis showed up-regulation of high numbers of genes in the cells exposed to low glucose, we also sought to characterize the changes in an activation mark, H3K4me3, induced by incubation of A549 cells in low glucose. We observed that a minority of H3K4me3 sites was changed by incubation in low glucose (Fig. 5F-G). As expected, most enrichment sites were associated with promoters or introns (Fig. 5H). Alignment of the ChIP-seq data with the RNA-seq data showed no significant enrichment of H3K4me3 on the promoters of genes that were activated in the RNA-seq experiment, suggesting that this histone mark is not associated with gene activation after 5 days of glucose starvation. However, we reasoned that changes in this activation mark could cause changes in gene expression at longer time points. We decided to focus on the promoters of the genes on which H3K4me3 mark is increased after incubation in low glucose. We identified 14 genes that displayed higher recruitment of H3K4me3 on their promoter in low vs high glucose (Table EV7). Transcription factor analysis of these promoters (Table EV8) showed a significant enrichment of binding sites for Snail, Slug, and ZEB1 in the promoters of these genes, suggesting that glucose restriction induces activation of epithelial-to-mesenchymal transition (EMT). Interestingly, the promoters of 12 of the 14 genes also contained the binding site for HIF-1α, confirming an important role of the hypoxia pathway in the response to glucose restriction. However, only 3 of the 14 genes were up-regulated by low glucose in the RNA-seq experiment. We reasoned that these genes may be primed for activation after 5 days of glucose restriction, but require longer incubation times to be up-regulated. Therefore, we performed RT-PCR analysis of the 14 genes in A549 cells incubated in low glucose for 5 days vs long-term incubation in low glucose for at least one month. As expected, 11 of the 14 genes were significantly (p-value: <0.01; t-test) up-regulated by longer incubation in low glucose (Fig.5I), whereas 3 did not result detectable in A549 cells by RT-PCR analysis. These observations suggested that glucose restriction primes certain genes associated with EMT transition for activation at later time points.

It has been previously reported that glutamine deprivation causes activation of Slug, but not Snail or ZEB1 (Recouvreux *et al*, 2020). To test the activation of Snail, Slug and ZEB1 in our system, we incubated A549 and NCI-H358 cells in high vs low glucose for 5 days, with or without supplementation of dm-αKG, followed by Western blot. Glucose deprivation caused increased level of Slug in both cell lines, rescued by dm-αKG, whereas it surprisingly caused a reduction in Snail and ZEB1 levels in A549 (this factor was not expressed in NCI-358 cells), and this inhibition was partially rescued by dm-αKG (Fig. 5J). This experiment suggests that Slug, but not ZEB or Snail, is up-regulated by low glucose, and it may be responsible for driving the more aggressive phenotype observed in glucose-restricted cells.

### 6. EZH2 causes cancer cell de-differentiation by regulating HIF1α signaling in LUAD cells

Since the hypoxia pathway was significantly up-regulated in the RNA-seq analysis, and HIF1α activation has been associated with EMT induction, we decided to focus on this pathway. To examine the activation of hypoxia-related pathways by glucose deprivation, we performed western blot analysis in A549 and NCI-H358 cells after incubation for 5 days in low glucose, with or without addition of dm-αKG. Both the two major HIF isoforms, HIF1α and HIF2α, were up-regulated by low glucose in A549 (Fig 6A) and NCI-H358 (Fig. EV4A) cells, and this was reversed by dm-αKG, whereas dm-αKG did not have any effect on the cells growing in low glucose. Furthermore, RT-PCR showed up-regulation of both HIF isoforms by low glucose, rescued by dm-αKG in A549 (Fig. 6B) and in NCI-H358 cells (Fig. EV4B). These results showed that low glucose can up-regulate the expression of HIF isoforms at both the protein and the mRNA level.

**Fig. 6.**
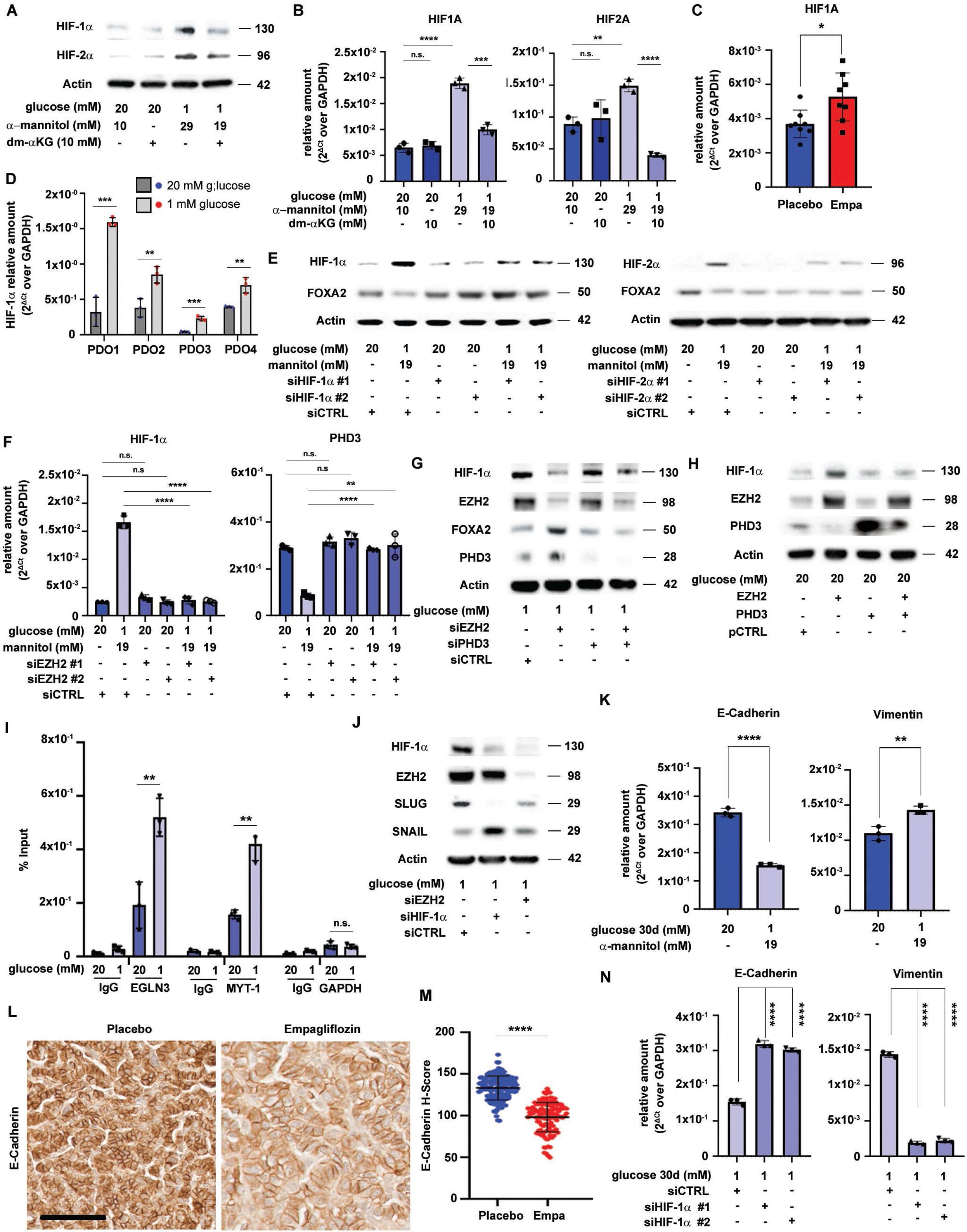
EZH2 causes de-differentiation by regulating Hif-1α signaling in LUAD. A-B) A549 cells were incubated in high or low glucose, with or without dm-αKG, as indicated. HIF1α and HIF2α expression was evaluated by western blotting (A) and RT-PCR (B). C) KP mice were give AdenoCre to induce tumors, followed by treatment with either placebo or empagliflozin as in Figure 1F, followed by collection of the lungs and RNA extraction for RT-PCR. Significance was measured by Student’s t-test. *p < 0.05. D) PDOs from 4 patients were incubated in high vs low glucose, as in Fig. 1I, followed by RT-PCR. Significance was measured by Student’s t-test. *p < 0.05; **P < 0.01; ***p < 0.001. E) A549 cells were incubated in low glucose and transfected with either control or siRNAs targeting HIF1α and HIF2α in either high or low glucose, followed by western blotting for HIF1α, HIF2α and FOXA2 to assess cell differentiation. F) HIF1A and PHD3 mRNA expression level was evaluated by RT-PCR in A549 cell transfected with or without siRNAs targeting EZH2. G) A549 cells were transfected with pooled siRNAs for EZH2 and/or PHD3. Knockdown efficiency was evaluated by western blotting, and changes in HIF1α and FOXA2 expression was evaluated by western blotting. H) HIF1α expression was evaluated in cells transfected with EZH2 and PHD3 expression vectors. The overexpression of the two proteins was also confirmed by western blotting. I) A549 cells were incubated in either high or low glucose for 5 days, followed by ChIP assay with either an antibody targeting EZH2 or normal IgG as negative control. qPCR was performed on the precipitated DNA with primers targeting the promoter of EGLN3 (encoding for PHD3), MYT-1 (a canonical EZH2 target) or GAPDH (an unrelated promoter as negative control). The results are reported as percentage of input. Significance was evaluated by Student’s T-test comparing the values in high vs low glucose for each target promoter. **p < 0.01; n.s. not significant. J) A549 cells were incubated for 5 days in low glucose, with transfection of either control siRNA or siRNAs targeting HIF1α and EZH2, followed by Western blot analysis as indicated. K) RT-PCR for E-cadherin and Vimentin was performed in A549 cells incubated in high glucose and low glucose for 30 days, and in cells incubated in low glucose for 30 days with transfection of siRNAs for HIF1α. Significance was measured by Student’s t-test. **p < 0.01; ****p < 0.0001. L-M) KP mice carrying LUADs were treated with either placebo or empagliflozin (10 mg/kg/d), starting 2 weeks after tumor induction for 6 weeks, as in Figure 1A. L) Representative images. M) Quantification of the E-Cadherin signal, expressed as H score, measured by QuPath analysis. Error bars: mean ± SEM. ****p < 0.0001. N) RT-PCR analysis E-Cadherin and Vimenin performed on A549 cells incubated in low glucose and transfected with siRNAs targeting HIF1α or siCTRL. Significance was measured by Student’s t-test. ****p < 0.0001

To confirm the activation of the hypoxia pathway in vivo and in more clinically relevant models, we performed RT-PCR for HIF1α in the KP murine tumors of mice treated with empagliflozin and in the four PDOs incubated in low glucose (the same experiments as in Fig. 1). As expected, both empagliflozin treatment of mice (Fig. 6C) and incubation of PDOs in low glucose (Fig. 6D) caused a significant up-regulation of HIF1α RNA levels.

To evaluate if HIFs were involved in glucose restriction-induced de-differentiation, we incubated A549 cells in low glucose with or without the transfection of siRNAs targeting either HIF1α or HIF2α. The efficiency of siRNA knockdown was also confirmed by western blot for both proteins. We assessed the effect of HIF knockdown on cell differentiation (FOXA2). Interestingly, we observed that the HIF1α, but not the HIF2α knockdown, increased the expression of LUAD differentiation marker FOXA2 in low glucose, but did not have any effect on the basal levels of FOXA2 in high glucose in A549 (Fig. 6E) and in NCI-H358 cells (Fig. EV4C), suggesting that activation of HIF1α repressed FOXA2 in low glucose. We next aimed to investigate the mechanism of HIF1α up-regulation by glucose restriction in LUAD. HIF1α is negatively regulated by prolyl hydroxylases (PHDs), which induce proteasome-dependent degradation of HIF1α under normoxic conditions (Xia *et al*, 2017). To investigate which PHD isoform was affected by low glucose, we performed RT-PCR for the three known PHDs in A549 and NCI-H358 cells incubated in high and low glucose with and without dm-αKG for 5 days. Only the expression of PHD3 was repressed by low glucose and rescued after the addition of dm-αKG in both A549 and NCI-H358 cells (Fig. EV4D). It has been previously demonstrated that loss of PHD3 promotes de-differentiation in breast cancer through a hydroxylase-dependent mechanism (Iriondo *et al*, 2015), and PHD3 is strongly up-regulated in well-differentiated human pancreatic cancer cells compared with less-differentiated tumors (Su *et al*, 2010), acting as a marker of differentiation. Consistently, PHD3 knockout wit htwo different siRNAs caused increased VEGF expression (a marker of HIF activation) and reduced FoxA2, suggesting de-differentiation in high glucose (Fig. EV4E). Therefore, we hypothesized that EZH2 up-regulates HIF1α and leads to de-differentiation in low glucose via inhibition of PHD3. To investigate the role of EZH2 in HIF1α regulation by glucose, we incubated cells in low glucose and performed transfection with either control or siRNAs targeting EZH2, followed by RT-PCR analysis. EZH2 knockdown reduced HIF1α and increased PHD3 expression in A549 (Fig. 6F) and in NCI-H358 cells (Fig. EV4F). This result was also confirmed by western blot: PHD3 was up-regulated and HIF1α down-regulated by EZH2 silencing in low glucose in A549 (Fig. 6G) and in NCI-H358 cells (Fig. EV4G). As expected, EZH2 siRNA also increased the expression of differentiation marker FOXA2, suggesting that EZH2 inhibits cell differentiation through PHD3 inhibition and HIF1α stabilization in low glucose. To verify this mechanism, we also transfected cells with an siRNA pool targeting PHD3. Both PHD3 siRNAs were previously tested separately in both cell lines (Fig. EV4H). PHD3 knockdown in low glucose did not cause significant changes in HIF1α and FOXA2 compared with control. However, double knockdown of both EZH2 and PHD3 protein resulted in increased expression of HIF1α and inhibition of FoxA2 compared with single transfection of siRNA for EZH2 (Fig 6G and EV4G). This result suggested that HIF1α activation and cell de-differentiation induced by EZH2 in low glucose are mediated by inhibition of PHD3. To confirm this mechanism of HIF1α stabilization, we over-expressed EZH2 and/or PHD3 in A549 cells incubated in regular media. EZH2 overexpression caused reduced PHD3 and increased HIF1α expression. PHD3 overexpression did not change the basal level of HIF1α, which was already basally low. Interestingly, the double overexpression of EZH2 and PHD3 abrogated the HIF1α induction caused by EZH2 overexpression, bringing back HIF1α to control levels in A549 (Fig. 6H) and in NCI-H358 cells (Fig. EV4H). These results suggest that the unbalanced activation of EZH2 in low glucose causes LUAD de-differentiation by inhibition of PHD3 and consequent activation of HIF1α. Interrogation of the Gene Transcription Regulation Database (GTRD) (Yevshin *et al*, 2017) showed recruitment of EZH2 on two sites in the promoter of the EGLN3 gene, which encodes for PHD3 (Fig. EV5A). To verify that EZH2 is directly responsible for transcriptional repression of PHD3 in low glucose, we performed ChIP-qPCR assay. We incubated A549 cells in either high or low glucose for 5 days, followed by ChIP with an antibody specific for EZH2. This experiment showed that EZH2 recruitment was increased in low glucose both on the EGLN3 promoter and on the canonical EZH2 target MYT-1, whereas no recruitment was observed on an unrelated promoter (GAPDH), or in the sample precipitated with control IgG (Fig. 6I). Similar results were obtained in NCI-H358 cells (Fig. EV5B). This result confirmed that EZH2 directly repressed the expression of PHD3 in cells incubated in low glucose.

Since the HIF pathway is associated with EMT and we have observed Slug induction by incubation in low glucose, we tested the hypothesis that Slug is downstream of the EZH2-HIF1α axis here identified. We incubated A549 cells in either high or low glucose, and in low glucose with control siRNA or pooled siRNAs targeting either EZH2 or HIF1α. As expected, incubation in low glucose caused increased levels of HIF1α and Slug, and reduction of Snail (Fig.6J). HIF1α knock-down completely rescued, and EZH2 knock-down partially rescued, the changes induced by low glucose in both Slug and Snail levels in A549 (Fig. 6J) and in NCI-H358 cells (Fig. EV5C). This experiment confirmed that Slug activation in low glucose is dependent on HIF1α. Our above-shown data showed that incubation for 5 days in low glucose primes lung cancer cells for EMT, but EMT-associated genes are up-regulated at later time points. Therefore, we aimed at investigating the up-regulation of Slug by long-term incubation in low glucose. We performed RT-PCR for Slug canonical inhibition target E-cadherin and the EMT marker vimentin after 30 days of low glucose incubation. In addition, we knocked down HIF1α in cells cultured in low glucose for 30 days. As expected, long incubation in low glucose repressed E-cadherin and increased vimentin expression in A549 (Fig. 6K) and in NCI-H358 cells (Fig. EV5D). The results showed that incubation in low glucose caused activation of Slug and subsequent EMT, with inhibition of E-cadherin and up-regulation of vimentin at 30 days. To confirm the relevance of this pathway *in vivo*, we performed IHC for E-cadherin in lungs of KP mice treated with empagliflozin for 6 weeks (same experiment as in Fig. 1A-E). As expected, SGLT2 inhibition caused a significant down-regulation of E-cadherin (Fig. 6L-M), confirming that this pathway is active *in vivo*. To confirm the role of HIF1α in low glucose-induced EMT, we incubated the cells in low glucose for at least 30 days, followed by transfection with siRNAs targeting HIF1α. As expected, E-cadherin expression was increased, and vimentin was reduced, by siHIF1α in A549 (Fig. 6N) and in NCI-H358 cells (Fig. EV5E). These data confirmed that pseudohypoxia in low glucose activates the EMT pathway in LUAD cells.

Our data suggested that the regulation of HIF1α by the EZH2-PHD3 pathway was transcriptional, since both RNA and protein levels of HIF1α were stimulated in low glucose. However, HIF1α is known to be regulated post-transcriptionally by PHD3, through prolyl-hydroxylation and proteasome degradation. Importantly, the activity of PHD3 also depends upon αKG availability. However, the direct hypoxic stabilization of HIF1α is an early event (within hours of hypoxia exposure), whereas the time point we examined here was after 5 days. After so long, more indirect mechanisms are likely to occur. In particular, it has been recently shown that increased HIF1α activity triggers a positive feedback loop with transcription of the long non-coding RNA HIFAL, which ultimately activates HIF1α transcriptionally (Zheng *et al*, 2021). We therefore measured the expression of HIFAL, and detected a significant activation of HIFAL in low glucose, completely rescued by αKG (Fig. EV5F-G). To verify that HIFAL is responsible for HIF1α induction after 5 days of low glucose exposure, we transfected A549 (Fig. EV5H) and NCI-H358 cells (Fig. EV5I) with siRNAs targeting HIFAL, and observed that HIFAL knockdown in low glucose causes reduction of HIF1α and GLUT1 expression and induction of FoxA2. These results suggest sustained HIF1α activation in low glucose exposure requires an HIFAL-dependent positive feedback with transcriptional up-regulation of HIF1α.

To confirm the hypothesis that short-term and long-term glucose deprivation cause activation of HIF1α through different mechanisms (protein stabilization and transcriptional regulation, respectively), we performed a time course experiment with incubation of A549 cells in low glucose for different time points, from 2 to 48 hours, showing that the increase in HIF1α protein levels is an early event, starting at 2 hours post-incubation in low glucose, and continuing for up to 48 hours (Fig. EV6A). We also explored earlier time points (30’, 1h and 2h) both in A549 and NCI-H358 cells, showing that 2h is the earliest time point when HIF1α is activated (Fig. EV6B). To verify the mechanism of early activation of the HIF1α pathway, we treated A549 and NCI-H358 cells both with αKG and with EZH2 knockdown and performed Western blot after 2 hours of incubation in low glucose. This experiment showed that after short-time glucose deprivation the activation of HIF1α is rescued by αKG, but not by EZH2 knockdown (Fig. EV6C). Taken together, these data suggest that glucose deprivation activates the hypoxia pathway through different mechanisms: at early time points (2h), HIF1α is regulated by αKG but not by EZH2, whereas at later time points (5d) HIF1α is regulated both by αKG and by EZH2. We assume that at 2h HIF1α stabilization depends on direct inactivation of PHDs by low αKG availability (Fig. 7A-B), whereas at later time points the activation of positive feedback loops including HIFAL up-regulation causes transcriptional up-regulation of HIF1α and EZH2 recruitment on the PHD3 promoter causes transcriptional repression of PHD3, with a combination of transcriptional and post-transcriptional activation of HIF1α (Fig. 7C).

**Fig. 7.**
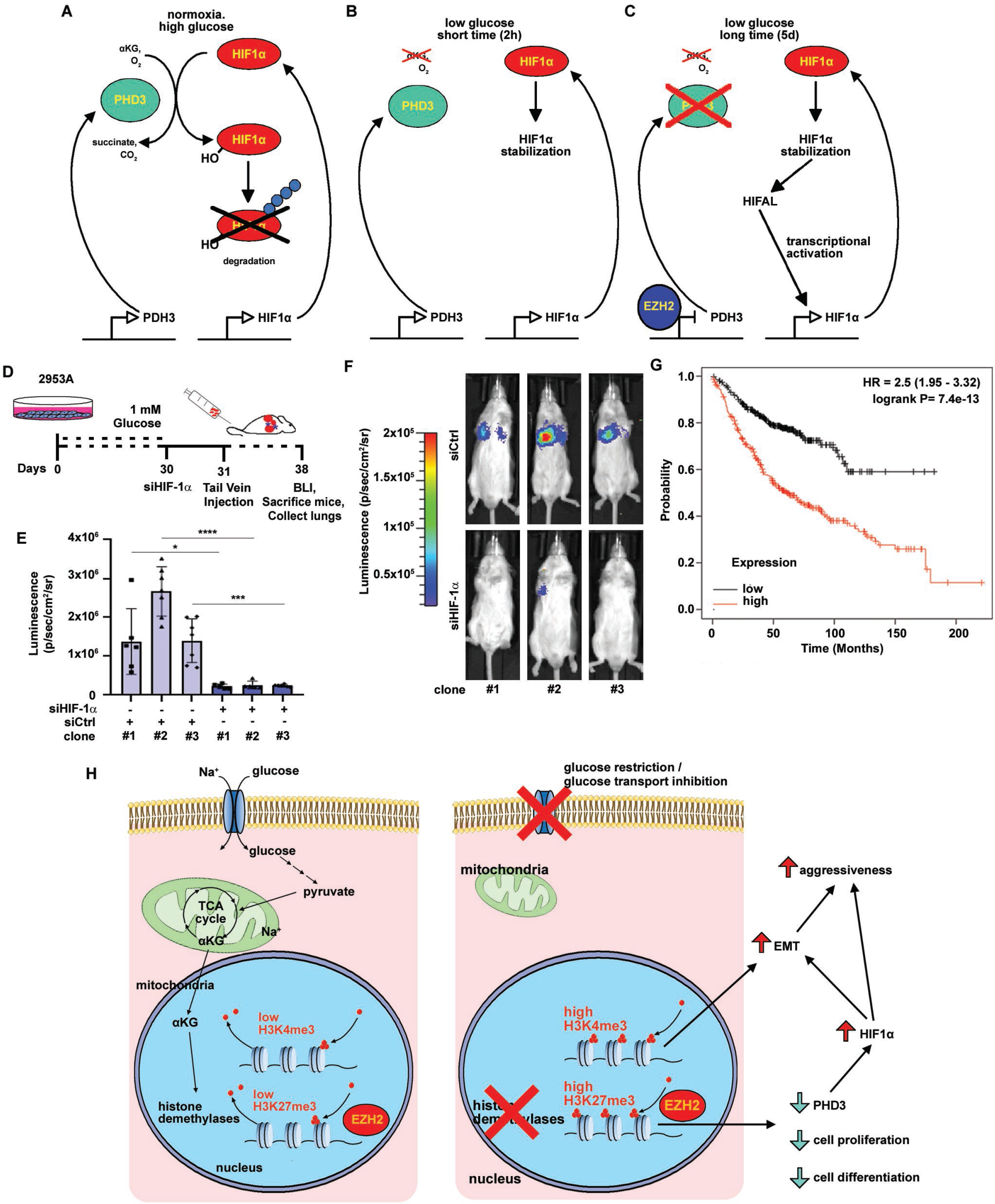
HIF-1α signaling is associated to a more aggressive phenotype. A-C) Model of HIF1α regulation by low glucose. In normal conditions of oxygen and glucose, HIF1α is targeted for degradation by αKG-dependent hydroxylation by PHD3 (A). When cells are incubated in low glucose for a short time (2h), the lack of αKG causes HIF1α stabilization, which can be rescued by αKG supplementation (B). Longer exposure to low glucose (5 days) causes more complex mechanism of HIF1α activation, with transcriptional up-regulation by the long noncoding RNA HIFAL, and transcriptional repression of PHD3 by EZH2 recruitment on the PHD3 gene promoter (C). D-F) Three different low glucose-induced clones of murine LUAD cells 2953A were expanded from a long-term incubation in low (1 mM) glucose for at least one month, as in Fig. 2D-F. One day before the injection in syngeneic mice, cells were transfected with two pooled siRNAs targeting HIF-1α and siRNA control. The day after, the cells were injected in syngeneic mice by tail vein injection to measure the development of lung metastases (D). The tumor burden was measured by bioluminescence. We report quantification of the bioluminescence (E) and representative bioluminescence pictures of single mice (F). Significance was evaluated in the pooled biological replicates by a general linear model with terms for group, experiment, and group x experiment interaction. *p < 0.05; ***p < 0.001; ****p < 0.0001. G) Kaplan-Meier survival curve of TCGA data showing a statistically significant difference in survival in patients expressing high vs low levels of genes involved concomitantly in HIF1A signaling.

We next investigated the role of HIF1α in the observed phenotypic shift towards a more aggressive phenotype. To this aim, we used murine 2953A cells incubated for at least 1 month in culture medium containing low glucose (1 mM). We transfected 3 clones of low-glucose 2953A cells with either control or pooled siRNAs against HIF1α, followed by tail vein inoculation in syngeneic mice (Fig. 7D). A control experiment with two different concentrations of the siRNAs targeting HIF1α showed that the RNAs were still silenced 7 days after transfection (Fig. EV6D). One week post-injection, we detected that the cells transfected with HIF1α siRNAs had significantly lower BLI signal than the control cells (Fig. 7E-F). These results suggest that HIF1α activation by glucose restriction is responsible for a transition of LUAD cells towards a more aggressive phenotype.

We finally sought to investigate the clinical relevance of the HIF pathway in human LUAD by interrogating The Cancer Genome Atlas (TCGA) database. We performed Kaplan-Meier analysis to measure the effect of hypoxia-regulated genes identified in our RNA-seq experiment on LUAD patient survival. Table EV9 shows the Hazard Ratio (HR) and p-value of hypoxia-regulated genes whose over-expression was associated with a significant HR (>1.5 or <0.5). These included, as expected, glycolytic genes (SLC2A1, HK2, ENO1, ENO2, ALDOC) and canonical HIF1α targets (VEGFA, VEGFB). We identified a group of 16 genes associated with HR >1.5 (ALDOC, ANGPTL4, CAVIN1, DDIT3, ENO1, ENO2, FAM162A, HK2, LDHA, PPP1R15A, PPP1R3C, SLC2A1, TPI1, VEGFA, VEGFB, VHL), and measured the cumulative effect of over-expression of these genes on LUAD patient survival. Cumulative expression of these genes was associated with an HR of 2.55 (p < 0.0001), confirming an important role of the activation of the hypoxia pathway on the clinical behavior of LUAD (Fig. 7G).

### 6. Nutrient restriction-induced de-differentiation is a general mechanism shared by different cancers

We finally aimed to check if glucose restriction-induced de-differentiation is restricted to lung cancer or generalized to other cancer types. Western blot analysis showed that incubation in low glucose for 5 days induced FoxA2 down-regulation, rescued by αKG, in a breast cancer (MCF7) and pancreas cancer (PANC1) cell line (Fig. EV6E). This result confirms that the described pathway is not restricted to lung adenocarcinoma, but is present in other tumors as well.

Glutamine depletion has been previously shown to cause H3K27 hypermethylation and de-differentiation in the core regions of melanoma (Pan *et al.*, 2016). We wanted to test if glutamine starvation also caused HIF1α activation in our system. We incubated A549 and NCI-H358 cells in either low glucose (1 mM) or low glutamine (2 mM), or both low glucose and glutamine, for 5 days, followed by Western blot analysis of HIF1α expression. As expected, glutamine deprivation caused an increase in HIF1α level, which was even more pronounced than that caused by glucose deprivation (Fig. EV6F). This is expected, because αKG derives directly from glutamine. Interestingly, restriction of both glutamine and glucose did not cause a further increase in HIF1α activation. Taken together, these data confirmed that pseudohypoxia is a common response mechanism to glutamine and glucose starvation.

## DISCUSSION

Here, we report a novel mechanism by which glucose restriction induces de-differentiation and increases aggressiveness of LUAD cells. Glucose deprivation causes reduced abundance of TCA cycle metabolites, with insufficient activity of αKG-dependent histone demethylases and unbalanced activity of methyltransferase EZH2, leading to repression of PHD3 and hyperactivation of HIF1α. HIF1α activation leads to Slug activation and EMT, eventually causing an aggressive/metastatic phenotype (Fig. 7H). We envision at least two different scenarios in which this mechanism can be relevant. The first is treatment with glucose transport inhibitors or glycolytic inhibitors, which are novel experimental strategies against lung cancer (Hsieh *et al.*, 2019; Scafoglio *et al.*, 2018). Our work provides evidence that these treatments are effective in reducing the tumor burden, but are also likely to cause an epigenetic adaptation of cancer cells to glucose restriction, leading to an unexpected and unintended de-differentiation and increased aggressiveness of treated tumors driven by pseudohypoxia. We showed that combination treatment with an EZH2 inhibitor, tazemetostat, significantly improves the response of LUAD to SGLT2 inhibition. Combination treatments with epigenetic modulators or HIF inhibitors are therefore important strategies as metabolic therapies are moved to the clinic.

In a different scenario, glucose deprivation can occur in cancers even in the absence of treatments, as a consequence of insufficient vascularization. In this case, glucose deprivation is accompanied by glutamine deprivation and hypoxia, reinforcing the activation of the HIF pathway and accelerating the progression of LUAD toward a more aggressive and de-differentiated phenotype. Consistently, our results showed that over-expression of certain genes of the hypoxia pathway in human LUAD is associated with significantly reduced survival.

Cancer de-differentiation is known to occur because of glutamine deprivation in the tumor microenvironment (Pan *et al.*, 2016). Our data suggested that glucose deprivation causes a similar phenotypic switch from early to poorly differentiated state in LUAD. We showed that αKG depletion causes an unbalanced activity between histone de-methylase and methyltransferase, resulting in hypermethylation on histone marks. In particular, increased tri-methylation on H3K27, carried out by EZH2, can affect the expression of a wide variety of signalling pathways involved in cellular transition to a more aggressive phenotype. Our work led to the discovery that a key mediator of the aggressive phenotype caused by glucose (and glutamine) deprivation is pseudohypoxia due to PHD3 inhibition by EZH2, and Slug activation.

Our ChIP-seq analysis showed that glucose deprivation causes a massive repositioning of EZH2-dependent modulation of histone of trimethylation of H3 lysine-27 in the genome, with predominance of effect in intergenic regions. The role of this histone mark on intergenic regions is only starting to be elucidated. Some of these intergenic loci are associated with silencer elements; the role of the majority of these intergenic sites is still to be discovered (Cai *et al.*, 2021). In addition, we observed that glucose depletion affects the tri-methylation in different histone marks, such as H3K4 and H3K9, as well as H3K27 acetylation. This suggests that several histone alterations can be potentially involved in the observed phenotypic transition. Enrichment of H3K4me3 on a subset of Slug target genes associated with EMT poises these genes for activation, promoting a more aggressive and metastatic phenotype. Further studies to dissect the role of different histone modifications and DNA methylation in glucose-induced cell de-differentiation are warranted.

We found that the HIF signalling pathway played a relevant role in glucose restriction-induced de-differentiation. This is very interesting because HIF1α can drive de-differentiation and cancer stem cell phenotype (Wang *et al*, 2017). Our data shows that EZH2 activation represses PHD3 expression due to direct repression with recruitment of EZH2 on the PHD3 promoter. In addition, PHD3 regulates the stability of HIF1α protein by inducing its ubiquitination and degradation. However, our data show induction of HIF1α both at the RNA and protein level, suggesting a transcriptional regulation of this gene by glucose restriction. HIF stabilization by hypoxia and by glucose deprivation occurs early within hours, without changes in HIF1α mRNA levels (Kallio *et al*, 1997; Wang *et al*, 1995). A short-term exposure to low glucose stabilized the HIF1α protein in an EZH2-independent manner, likely because the PHD proteins responsible for targeting HIFs for degradation are αKG-dependent enzymes (Jaakkola *et al*, 2001; Kivirikko & Myllyharju, 1998). Long-term regulation of HIF1α involves more complex combinations of transcriptional and post-transcriptional mechanisms, with involvement of feed-forward loops and long non-coding RNA-mediated regulation (Zheng *et al.*, 2021). EZH2-mediated regulation of HIF1α observed in our studies is likely involved in long-term regulation of HIF1α expression, linking glucose starvation with changes in cell differentiation state.

In conclusion, we discovered a novel mechanism of glucose restriction-induced cancer de-differentiation, mediated by unbalanced activity of histone methyltransferase EZH2 and activation of the hypoxia pathway and EMT, leading to an aggressive phenotype. These results should be considered in designing new anti-cancer metabolic therapies targeting glucose uptake and glycolysis.

## METHODS AND PROTOCOLS

### EXPERIMENTAL MODELS

#### Mouse models

All experiments performed in mice were approved by the UCLA Institutional Animal Care and Use Committee and were carried out according to the guidelines of the Department of Laboratory Animal Medicine (DLAM) at UCLA. For our imaging and therapeutic trials in genetically engineered mouse models (GEMMs) we used Kras^*LSL/*G12D^, p53^*fl/fl*^, Rosa26^*LSL/luciferase*^ mice (KPluc mice, in FVB background) bred in our laboratory, as previously reported (Scafoglio *et al.*, 2018). The breeders were kindly provided by Dr. David Shackelford (UCLA).

For tail vein injection assays we used syngeneic wild type mice (FVB background). KPluc mice were bred in our colony at UCLA. Syngeneic wild type mice were purchased from Jackson lab.

#### Cell lines

Human lung cancer A549 (CCL-185) and NCI-H358 (CRL-5807) cells lines were purchased from American Type Culture Collection (ATCC, Manassas, VA). Cells were maintained in culture according to manufacturer protocol using Roswell Park Memorial Institute (RPMI) 1640 medium (Corning, Cat. #10-040CV) supplemented with 10% FBS and 5% 1X Pen-Strep and kept in 37°C and 5% of CO_2._ Murine 2953A cell line was established in our lab from a KP lung tumor by tissue dissociation. This cell line was maintained in culture in Dulbecco's modified Eagle Medium (DMEM) medium (Corning, Cat. #10-017CV) with 5% FBS and 5% 1X Pen-Strep.

#### Patient-derived organoids culture

Patient-derived organoid (PDOs) lines were established in our lab from fresh human lung tumor procured from surgical pathology after obtaining informed consent (UCLA IRB #10-001096, PI: Steven Dubinett). The tissues were minced with surgical scissors into small (<0.5 mm) fragments and digested in lung dissociation mix (1500 μg/mL Collagenase A, 100 μg/mL Collagenase Type IV, 100 μg/mL DNase I, 100 μg/mL Dispase II, 9.2 μg/mL Elastase, 1250 μg/mL Pronase in Hank’s Balanced Salt Solution) for 2 cycles of 10 minute incubation on orbital shaker followed by mechanic dissociation by pipetting. When the tissue was completely dissolved, single cells were filtered through 70 μm cell strainer (Falcon). The enzymes were inactivated with FBS (final 3%). Pellets were resuspended first in ACK buffer (Gibco) for erythrocyte lysis and next in Advanced DMEM/F12 (1x Glutamax, 10 mM HEPES, 1X Anti-Anti) to stop the reaction. Cells were counted and seeded at 75000 cells per 50 μl of Cultrex Basement Membrane Extract (RGF BME; R&D Systems) and cultured in DMEM/F12 (ThermoFisher 11320033) supplemented with R-Spondin 1 (Peprotech #120-38, 500 ng/ml), FGF7 (Peprotech #100-19, 25 ng/ml), FGF10 (Peprotech 100-26, 100 ng/ml), noggin (Peprotech 120-10C, 100 ng/ml), A83-01 (Tocris #2939, 500 mM), Y-27632 (Abole #Y-27632, 5 μM), SB202190 (Sigma #S7067, 500 nM), B27 supplement (gibco #17504-44, 1x), N-acetyl-cysteine (Sigma #A9165-5g, 1.25 mM), nicotinamide (Sigma #N0636, 5 mM), glutamax (Invitrogen #12634-034, 1x), Hepes (Invitrogen #15630-056, 10 mM), penicillin/streptomycin (Invitrogen #15140-122, 100 U/ml and 100 μg/ml, respectively), primocin (Invivogen #Ant-pm-1, 50 μg/ml).

### PROTOCOLS

#### In vitro studies

All experiments in cell lines were performed in biological triplicate. Cells were seeded at same confluence and grown for 5 days (and for some experiments overnight or up to 9 days or 30 days, as indicated in the figures and text) in different concentration of glucose: high (20 mM), low (1 mM) and for some experiments medium (5 mM), as indicated. We used mediums with glutamine and without glucose: RPMI (Corning Cat. #10-043CV) for A549 and NCI-H358 cells, DMEM (ThermoScientific Cat. #11566025) for 2953A cells, supplemented with 10% FBS, 5% of 1X Pen-Strep and complemented with D-Glucose to the desired final concentration, as indicated in the figures.

For the glutamine deprivation experiment, A549 and NCI-H358 cell lines were also seeded and grown up to 5 days in DMEM without glutamine and glucose (Corning Cat. #17-207CV) and complemented with L-glutamine (GlutaMAX CTS, Gibco) to a final concentration of either 6 mM (high glutamine) or 2 mM (low glutamine) and D-Glucose to a final concentration of either 10 mM (high glucose) or 1 mM (low glucose). Finally, breast cancer (MCF7) and pancreatic cancer (PANC1) cell lines were purchased from ATCC and were seeded and grown up to 5 days in RPMI1640 complemented with D-Glucose to a final concentration of either 20 mM or 1 mM.

All αKG rescue experiments were conducted using 10 mM alpha-ketoglutarate (α-KG; Sigma) and α-Mannitol (Sigma) as osmotic control.

PDO lines were cultured for 3 weeks in Advanced DMEM/F12 without glucose (ThermoFisher #A2494301) complemented as described above for PDO culture, and with D-Glucose to a final concentration of either 20 mM (high) or 5 mM (medium) for immunofluorescence, and either 20 mM (high) or 1 mM (low) glucose for RNA extraction and RT-PCR. α-Mannitol was used as osmotic control In PDOs incubated in low glucose.

#### In vivo studies

Lung tumors were induced by intranasal administration of Adeno-Cre (purchased from University of Iowa Viral Vector Core) as previously described (Scafoglio *et al.*, 2018), or by transthoracic injection of Adeno-Cre in KP_luc_ mice. All treatment trials were performed 2 weeks after tumor induction, and the results of biological duplicates were pooled for statistical analysis. In all trials, mice were assigned to therapeutic groups so that there were no significant differences in baseline tumor burden, age, sex, or weight among the different groups.

In the trial for assessment of tumor differentiation by immunohistochemistry (Figure 1A-E, 6L-M, and EV1A-C) lung tumors were induced by intranasal administration of Adeno-Cre. After tumor induction, mice (n = 11 for placebo and 13 for empagliflozin group) were treated with either placebo (DMSO 5%, PEG 30%, Tween80 10%) or empagliflozin (10 mg/Kg/d by oral gavage) for 6 weeks and then sacrificed for lung collection and fixation in formalin.

In the trial for assessment of tumor differentiation by RT-PCR (Figure 1F and 6C) mice (n = 3) received intranasal AdenoCre and treatment with either placebo (0.5% hypromellose-methyl cellulose) or empagliflozin (10 mg/Kg/d by oral gavage) for 4 weeks, followed by tumor collection and RNA extraction. RT-PCR was performed in technical triplicates, yielding a total of 9 experimental points per group.

For the measurement of in vivo HeK27 trimethylation (Fig. 3B), mice (n = 4 per group) received transthoracic adenoCre and were treated for 3 weeks with either placebo (0.5% hypromellose-methyl cellulose) or empagliflozin (10 mg/Kg/d by oral gavage) followed by lung collection and histone extraction for ELISA assay.

In the combination treatment trial (Fig. 3F-G), mice received the first BLI measurement two weeks post-AdenoCre administration for measurement of the baseline tumor burden. Four therapeutic groups were established as follows: (i) Placebo (n = 11), receiving oral gavage of vehicle (DMSO 5%, PEG 30%, Tween80 10%); (ii) SGLT2 inhibitor Empaglifozin (10 mg/Kg/d, n = 13); (iii) EZH2 inhibitor Tazemetostat (125 mg/Kg/bid, n = 10); (iiii) Empaglifozin and Tazemetosat (n = 19). Each group received bi-daily administration by oral gavage.

#### Tail vein injection assays

For our studies on the investigation of cell aggressiveness due to the glucose deprivation, we used syngeneic FVB mice. Two experiments were performed, in biological duplicate.

In the first experiment (Fig. 2D-F and 7D-F), we used the murine cell line, 2953A, previously established in our lab from a KP tumor, which has a transgenic luciferase expression. 2953A cells were cultured in either high (20 mM) or low (1 mM) glucose for at least 1 month. In cells incubated in low glucose, α-mannitol was used as osmotic control. For low-glucose cells, to exclude clonal effects, we expanded three different clones of low glucose-resistant cells. The day before the inoculation, cells were transfected with either pooled siRNAs targeting HIF-1α or control siRNA. Cells were trypsinized and resuspended in cold PBS, followed by tail vein inoculation (250,000 cells/mouse) in syngeneic mice (n = 8). After 1 week, tumor burden was measured by bioluminescence performed on an IVIS Spectrum In Vivo Imaging System (PerkinElmer) 10 min after intraperitoneal injection of luciferin (150 mg/kg). Bioluminescence acquisitions were analyzed by Living Image Software (Perkin Elmer). For simplicity, we presented the high vs low glucose results in figure 2D-F and the control vs siHIF1α in low glucose in figure 7D-E.

In the second experiment (Fig. 2G-J), KP mice injected transthoracically with AdenoCre were treated for 3 weeks with either placebo (n = 3) or empagliflozin (n = 3), and sacrificed for lung collection. Tumors were identified in the murine lungs by ex vivo bioluminescence and dissociated. Tumor single cells were resuspended in cold PBS followed by magnetic beads sorting (Miltenyi Biotec) to isolate the Epcam-positive epithelial cells. Finally, sorted tumor cells were injected by tail vein inoculation (1,000,000 cells/mouse) into syngeneic mice (n = 6 per group). Tumor burden was measured by bioluminescence until 13 weeks after inoculation.

#### Immunohistochemistry (IHC) staining

For the GEMMs, the mouse lungs were collected and inflated with 10% formalin in phosphate-buffered saline. After 24 hours, formalin was replaced to 70% ethanol. All tissues were paraffin-embedded and sliced into 4-mm sections in the Translational Pathology Core Laboratory (TPCL) at UCLA.

For IHC staining, the slides were deparaffinized by overnight incubation at 65°C, followed by rehydration by serial passages in xylenes (three washes of 5min in 100% xylenes) and decreasing concentrations of ethanol (two washes in 100% ethanol, two washes in 95%, one wash in 80%, one wash in 70%, and one wash in water). Antigen retrieval was performed for 23 min in AR6 (pH 6.0) buffer.

Blocking was performed with 5% goat serum for 1 hour at room temperature, followed by incubation with primary antibodies overnight at 4°C. Incubation with biotin-labeled secondary antibody was performed at room temperature for 1 hour, followed by incubation with avidin-biotin peroxidase complex and ImmPACT 3,3′-diaminobenzidine (DAB). Counterstain was performed with Harris’ haematoxylin diluted 1:5 in water, followed by rehydration step and mounting slide. After the staining, digital images of the slides were obtained with an Aperio ScanScope slide scanner (Leica Biosystems). The quantification was performed using the QPath software. Briefly, regions of interests were drawn blindly in each slide to include each tumor present in a whole-lung section, followed by cell detection and quantification of the DAB signal using a constant threshold for all samples in the same experiment. Staining intensity in each cell was classified as negative (0), weak (1), median (2), or strong (3). The results were expressed as H score.

#### Immunofluorescence (IF) staining

PDOs were fixed in formalin and embedded in histogel and paraffin for sectioning. For immunofluorescence, the slides were de-paraffinized in xylene and serial dilutions of ethanol (as for IHC), followed by antigen retrieval in a vegetable steamer for 20’, washess, and sequential incubation in primary antibodies (HTII-280 and panCK), secondary antibodies (IgM-HRP), and Vectra Polaris chromophores Opal 520 (for HTII-280) and Opal 690 (for panCK). After washes, the slides were cover-slipped with mounting medium containing 300 nM DAPI. Images were acquired with Akoya Vectra Polaris slide scanner and analyzed with QPath software, as described above.

#### Metabolomics assay

A549 and NCI-H358 cells were cultured overnight in medium complemented with either high (20 mM) or low glucose (1 mM). For each condition, we prepared 3 plates for LC-MS analysis. Then, cells were rinsed with ice-cold 150 mM NH4AcO at pH 7 and incubated with precooled 80% methanol in −80°C for 60 minutes. Next, cells were scraped, transferred into a new tube on ice and spun down at 16,000g for 15 min at 4°C. Afterward, the supernatants, containing the metabolites extracted, were transferred in a glass vial and then they were dried down at 30°C in an evaporator (Genevac EZ-2 Elite). The pellets, which are still in the tubes on ice, were resuspended with 3 volumes of RIPA buffer and protein concentration was determined by BCA assay.

LC-MS was performed by UCLA Metabolomic Center by using the Q Exactive mass spectrometers coupled to an UltiMate 3000 UPLC chromatography systems (Thermo Fisher Scientific). Samples were normalized by protein content. Data collected was processed with TraceFinder 4.1 (Thermo Fisher Scientific). Histograms were generated considering on normalized relative amounts of each replicate of both groups. Statistical analysis was performed by analysis of variance (ANOVA).

#### Small interfering RNA transfection

For small interfering RNA (siRNA) knockdown of human EZH2, HIF-1α, HIF-2α and PHD3 cells were culture either high (20 mM) and low glucose (1 mM) and transfected with 25 pM of siRNAs for each target and control siRNA using RNAi Max reagent (Thermo Fisher scientific).

Since the total incubation time in low glucose was 5 days, two separate transfections were performed at day 1 before starting the high vs low glucose incubation, and at day 3. At day 5, cells were collected, followed by total protein extraction or RNA extraction. Each siRNA was used individually or pooled. Each pooled siRNA were transfected either as a single pool or in combination with another pooled siRNA.

For the long-term (30 days) exposure to low glucose (Fig. 6N and EV5E), cells were incubated long-term without transfection, and siRNAs were transfected 5 days and re-transfected 3 days before harvesting. For tail vein injection, cell clones isolated from long term low-glucose culture were transfected with a pool of two siRNAs targeting HIF-1α or control the day before tail vein injection. We confirmed the knockdown efficiency of mouse HIF-1α knockout 3 and 7 days after RNA transfection (Fig. EV6D).

All siRNAs were purchased from QIAGEN and Dharmacon (See Reagents and Tools table and Table EV10).

#### Plasmid transfection

A549 cell line was transfected with 10 μg of EZH2 and PHD3 plasmid, purchased from Addgene, using Lipofectamine 3000. Each plasmid was transfected either individually or in combination. After three days, cells were harvested, lysed with RIPA buffer and analysed by SDS-PAGE and Western blotting.

#### Enzyme-linked immunosorbent assay (ELISA)

The levels of trimethylation on H3K27me3 in tumors treated with empaglifozin were detected by ELISA kits (Active Motif) according to the manufacturer's instructions. The experiment was carried out with 4 replicates for each group. After tumor collecting, histone extraction was performed.

We used H3K27me3 and H3 total antibodies for plate coating. The absorbance was detected by Varioskan Lux (Thermo Fisher Scientific). The H3K27me3 levels were normalized to H3 total and presented as optical densitometry (O. D.) 450 nm of a ratio of H3K27me3/H3.

#### Total Protein Extraction

For total protein extraction, cells were harvested, washed twice with ice-cold PBS-EDTA (0.5 mm EDTA), lysed using RIPA buffer (50 mM Tris-HCl pH 7.6, 150 mM NaCl, 0.1% SDS, 0.5% C24H39NaO4, 1% NP-40, 2 mm EDTA, 50 mm NaF) for 15 min on ice and centrifuged at 13,000 rpm for 30 min at +4 °C. The resulting protein extracts were quantified using BCA protein assay followed by analysis by SDS-PAGE and Western blotting.

#### Histone extraction

Histones was extracted from A549, NCI-H358 and 2953A cell lines using the Histone extraction kit (Abcam) according to manufacturer instructions.

#### Western Blotting

SDS-PAGE and Western blotting analyses were performed using standard protocols. For antibodies information, see Reagents and Tools Table. Western blot images were detected by iBright Imaging Systems (Thermo Fisher Scientific). The protein detected was normalized to Actin or H3 total antibodies, this last in case of histonic extract. Each immunodetection derived from the same membrane was performed with the same exposure times according to the manufacture’s antibody guidelines and was cropped only for presentation purposes; this is indicated by a dotted line.

#### RNA extraction

Total RNA was extracted from A549, NCI-H358 and 2953A by using TRI Reagent Solution (Applied Biosystem), according to manufacturer instruction. RNA concertation was assayed by Nanodrop 3000 spectrophotometer (Thermo Fisher Scientific). Then, 1μg of RNA was treated with DNase I (Thermo Fisher Scientific) and used for cDNA preparation.

#### RT-qPCR

cDNA was prepared using 1 μg of RNA with SuperScript IV Reverse Transcriptase (Invitrogen). SYBR green-based RT-PCR kit (Biorad) was performed using human and mouse primers (See Table EV11). mRNA levels were normalized to GAPDH (ΔCt=Ctgene of interest−Ct GAPDH) and presented as relative mRNA expression (ΔΔCt = 2−(ΔCtsample–ΔCtcontrol)). All primers were designed by NCBI Primer-BLAST and purchased from Integrated DNA Technologies.

#### RNA-seq and Data Analysis

A549 and NCI-H358 cell lines were incubated in triplicate with high glucose (20 mM), low glucose (1 mM), and low glucose + dm-αKG (10 mM). Total RNA was extracted as previously described. RNA sequencing was performed by Med Genome. Sample quality control was performed using Quibit fluorometric (Agilent) and Tapestation bioanalyzer (Agilent). For library preparation TruSeq Stranded Total RNA kit (Illumina) was used and libraries were sequenced on NovaSeq platform (Illumina). The FASTQ data generated was used for gene expression analysis, performed as described in Nassa et al(Nassa *et al*, 2019).

Briefly, the raw sequence files generated (.fastq files) underwent quality control analysis using FASTQC (http://www.bioinformatics.babraham.ac.uk/projects/fastqc/) and adapter sequences were removed using Trimmomatic version 0.38(Bolger *et al*, 2014) Filtered reads were aligned on human genome (assembly hg38) considering genes present in GenCode Release 35 (GRCh38.p12) using STAR v2.7.6a(Dobin *et al*, 2013) with standard parameters. Quantification of expressed genes was performed using featureCounts(Liao *et al*, 2014) and differentially expressed genes were identified using DESeq2(Love *et al*, 2014). A given RNA was considered expressed when detected by at least ≥10 raw reads.

Differential expression was reported as |fold-change| (FC) ≥1.5 along with associated adjusted pvalue≤0.05 computed according to Benjamini-Hochberg. Gene Set Enrichment Analysis (GSEA) was performed to examine pathway enrichment for the differential expressed genes with the Molecular Signature Database “Hallmarks” gene set collection(Liberzon *et al*, 2015). Only those with an FDR ≤0.25 have been selected.

Functional analysis was also performed with Ingenuity Pathway analysis (IPA, QIAGEN). Only pathways with a pvalue ≤0.05 were considered for further analysis.

#### ChIP-seq and Data Analysis

A549 cell lines were incubated in triplicate with high glucose (20 mM) and low glucose (1 mM) for 5 days. A total of 15 ×10^6^ cells were fixed, lysed to isolate nuclei, sonicated, and diluted as described by Nassa et al. (Nassa *et al.*, 2019). An aliquot of nuclear extract was taken as input to be used as control for sequencing and data analysis. For H3K27me3 and H3K4me3 pull-down, 50 μl of equilibrated Dynabeads M-280 Sheep Anti-Rabbit IgG (Thermo Fisher Scientific) were incubated overnight at 4°C with 10 μg of the antibodies chosen for immunoprecipitations (Abcam). Bead washing, elution, reverse crosslinking and DNA extraction were then performed as described in Tarallo et al. (Tarallo *et al*, 2017). Size distribution of each ChIP DNA sample was assessed by running a 1 μl aliquot on an Agilent High Sensitivity DNA chip using an Agilent Technologies TapeStation (Agilent Technologies). The concentration of each DNA sample was determined by using a Quant-IT DNA Assay Kit-High Sensitivity and a Qubit Fluorometer (Life Technologies). Purified ChIP and input DNAs (6 ng each) were used as the starting material for sequencing library preparation by using the TruSeq ChIP Sample Prep Kit (Illumina Inc.) and were sequenced (single read, 1 × 75 cycles) on a NextSeq 500 (Illumina Inc.).

Quality control of the sequenced reads was performed using FASTQC (http://www.bioinformatics.babraham.ac.uk/projects/fastqc/). Reads were aligned to the reference genome assembly (hg38) using bowtie (Langmead *et al*, 2009) allowing up to one mismatch and considering uniquely mapped reads. Signal artefact blacklist regions were filtered out using bedtools (Quinlan & Hall, 2010).

For each biological replicate, peak calling was performed using HOMER (Heinz *et al*, 2010) setting the following parameter: −P 0.01 −F 2.5 −style histone. Only peaks common to at least two replicates were considered for further analysis. Annotation of peaks to the nearest gene was performed using the annotatePeaks.pl function of HOMER, while annotation of peaks to cis-Regulatory Elements was performed with bedtools, using the track available in SCREEN (Moore E et al., Nature, 2020).

To find differentially regulated features between low glucose versus high glucose the getDifferentialExpression.pl function of HOMER was used, setting the parameter “–edgeR”. Only features showing an adjusted p-value ≤0.05 and |FC|≥1.3.

Prediction of potential transcription factor binding sites on selected genes associated to histone peak was performed using Ciider (Gearing *et al*, 2019).

#### ChIP-qPCR

A549 cell line was incubated in either high (20 mM) or low (1 mM) glucose for 5 days. Chromatin was isolated as described previously starting from 15×10^6^ cells. Before immunoprecipitation, an aliquot of chromatin extract was taken as input to be used as control of qPCR. ChIP was carried out by over-night incubation of chromatin at 4°C with 50 μl of Dynabeads Protein G (Invitrogen), pre-coated with 5 μg of anti-EZH2 (CST). Beads washing steps, DNA elution and extraction were performed as previously described.

Before DNA elution, an aliquot of beads for each condition was conserved for western blot assay, resuspended in sample buffer and boiled at 90°C for 5 minutes.

0.2 ng of DNA were used to amplify PHD3, MYT-1 and GAPDH promoter regions (See Table EV12). Data analyses were represented as percentage of Input. The experiment was repeated a second time and the data were pooled as a biological replicate.

### QUANTIFICATION AND STATISTICAL ANALYSIS

Prism 8.0 software (GraphPad) was used for statistical analysis. Analysis for significance was performed by parametric or nonparametric Student t-test when only two groups were compared and by one-way ANOVA when three or more groups were compared.

For the treatment trials, in order to compare tumor volume (log scale) between groups (high glucose vs low glucose, control siRNA vs HIF1α siRNA), we ran a general linear model with terms for group, experiment, and group x experiment interaction. We then extracted the pairwise contrast from the model to compare the groups with 95% confidence intervals. P-values < 0.05 were considered statistically significant and all analyses were run using IBM SPSS V27 (Armonk, NY).

QuPath (Bankhead *et al*, 2017) was used for signal quantification in IHC staining. It was expressed as H score. Kaplan-Meier curves were performed with Kaplan-Meir Plotter tool (Gyorffy, 2021), using the lung cancer section. For the selected genes only, the best probe was used. All dataset available were used as cohort. For Kaplan-Meier plot performed using multiple genes, the option “use mean expression of selected genes” was set.

### RESOURCE AVAILABILITY

#### Lead contact

Further information and requests for resources and reagents should be directed to and will be fulfilled by the lead contact, Claudio Scafoglio.

#### Materials availability

This study did not generate new unique reagents.

#### Data and code availability

The RNA-seq raw data are publicly available in ArrayExpress repository under accession number: E-MTAB-11253.

The ChIP-seq raw data are publicly available in ArrayExpress repository under accession number: E-MTAB-11678.

Any additional information required to re-analyze the data reported in this paper is available from the lead contact upon request.

## ACKNOWLEDGEMENTS

This research was supported by the following grants: American Cancer Society grant number 130696-RSG-17-003-01-CCE (Scafoglio), NIH/NCI R01CA237401-01A1 (Scafoglio), UCLA Jonsson Comprehensive Cancer Center Seed Grant (Scafoglio), Fondazione AIRC, grant IG-23068 (Weisz) and Regione Campania POR Campania FESR 2014/2020 - Azione 1.5 - CUP: B41C17000080007, grant GENOMAeSALUTE (Weisz). PS was supported by an Italian American Cancer Foundation Fellowship.

We thank Dr. David B. Shackelford for providing the KP breeder mice to start the colony used for this project, and the UCLA Metabolomics Core for the metabolic LC-MS analyses.

## Author Contributions

CS and PS developed the study concept and experiment design. PS and AP performed all the experiments with assistance from AH and EF. ER performed pathologic analysis of murine tumors. TG provided statistical support. GG and AW performed RNA-seq data analyses and interpretation. AS and GN performed ChIP-seq experiments. AS, GG, GN and AW performed ChIP-seq data analyses and interpretations. SMD provided advice throughout the development of the project. PS and CS wrote the manuscript with input from all the authors.

## Declaration of Interests

The authors declare no conflicts of interest.

---

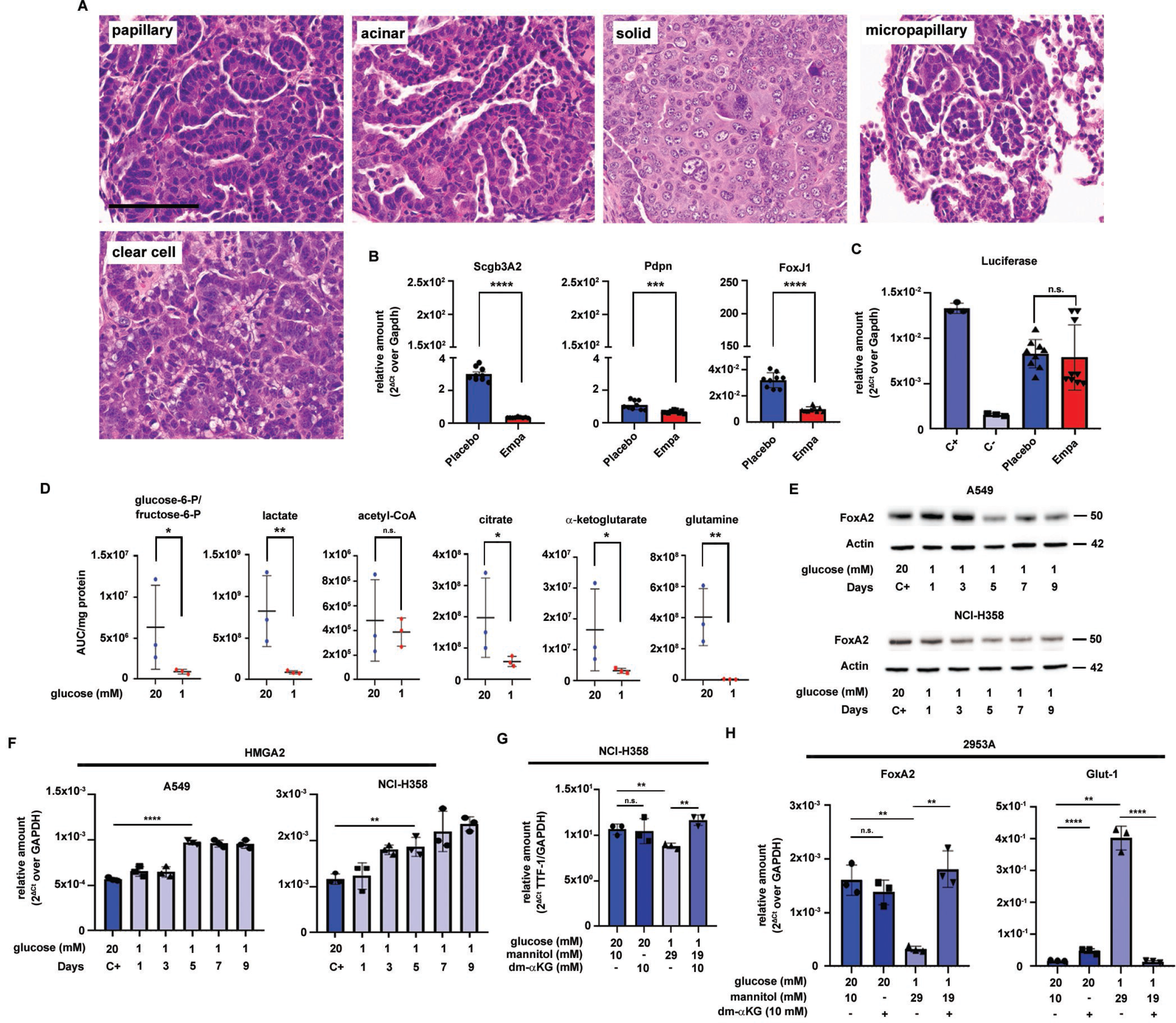

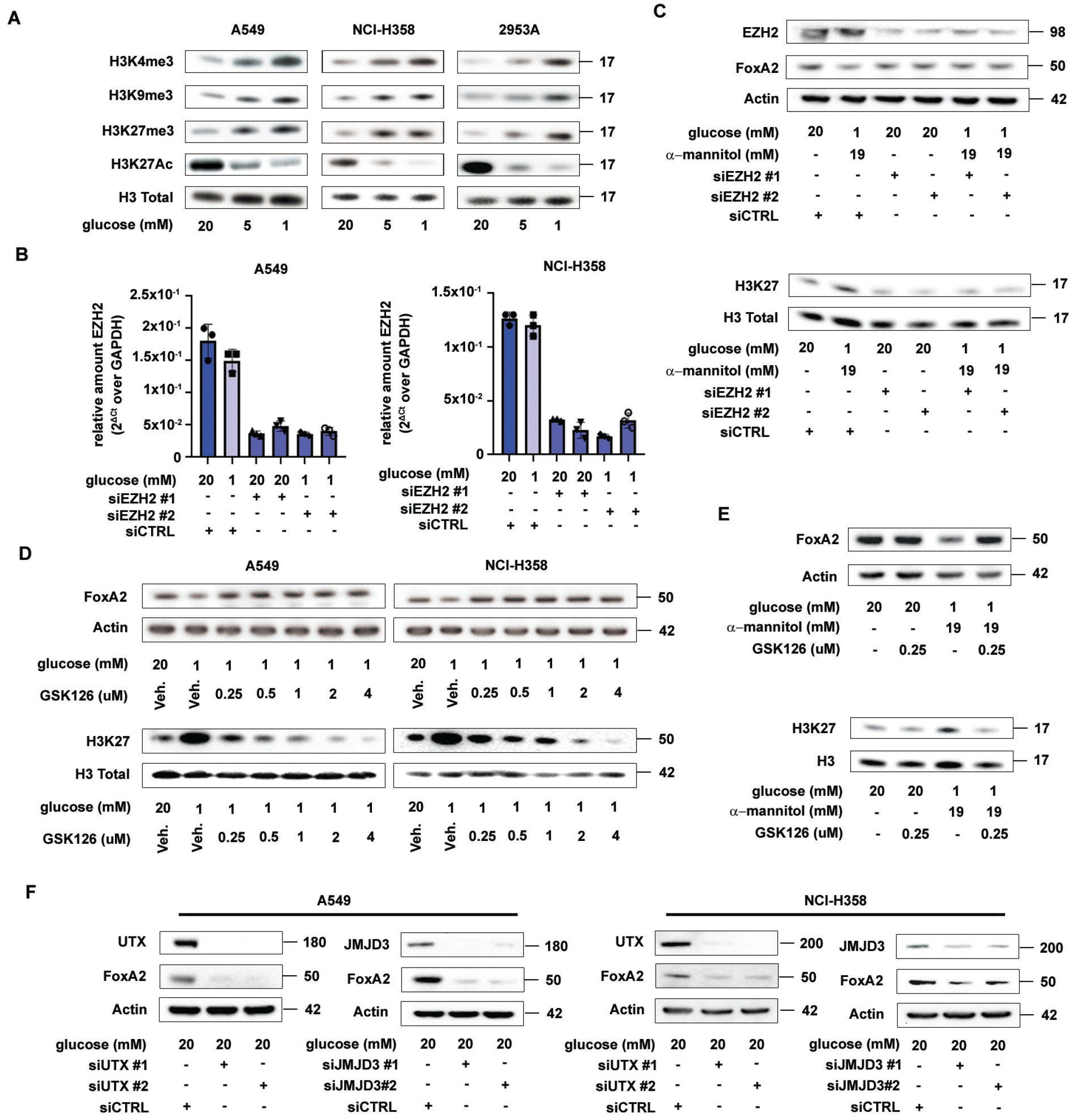

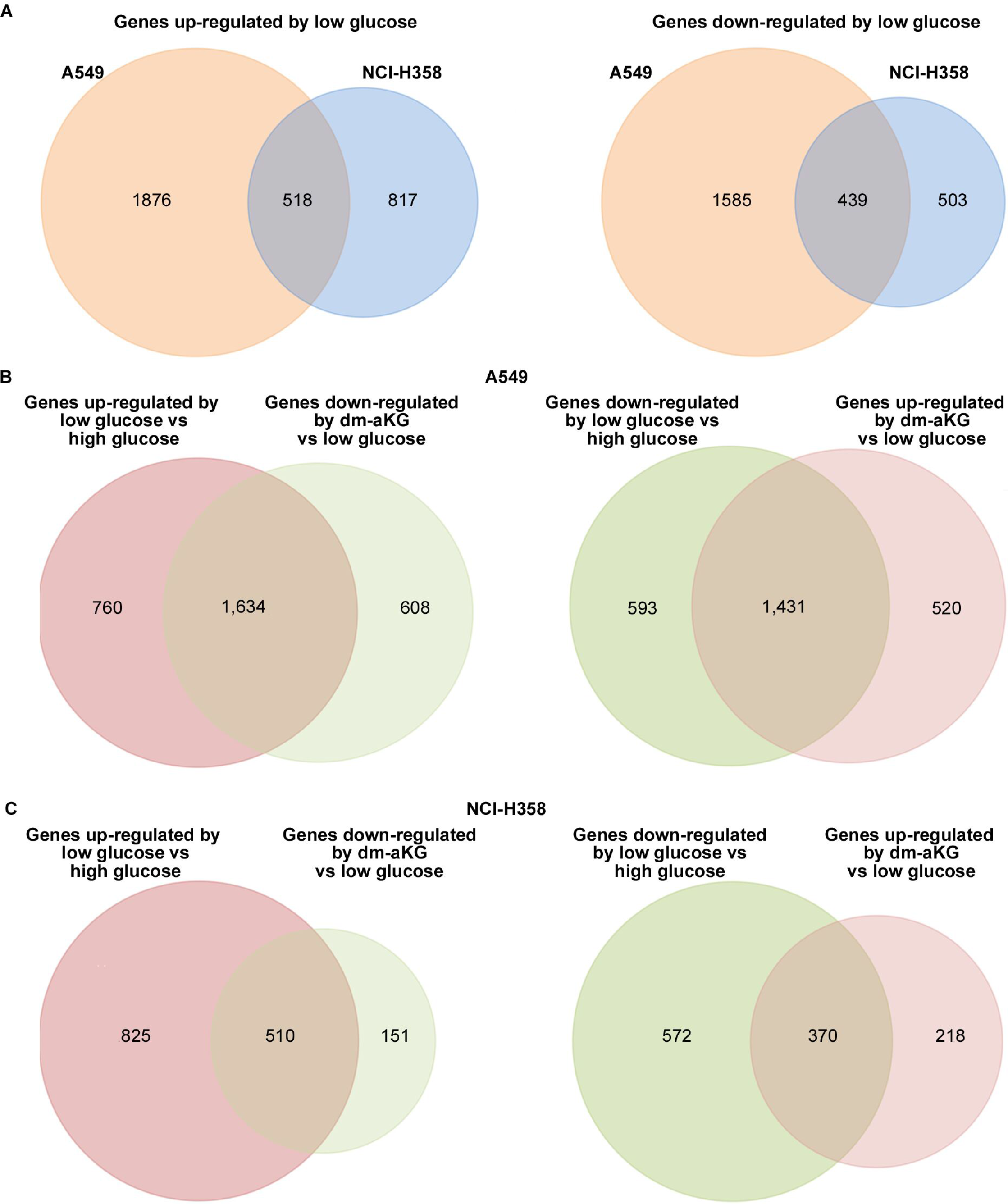

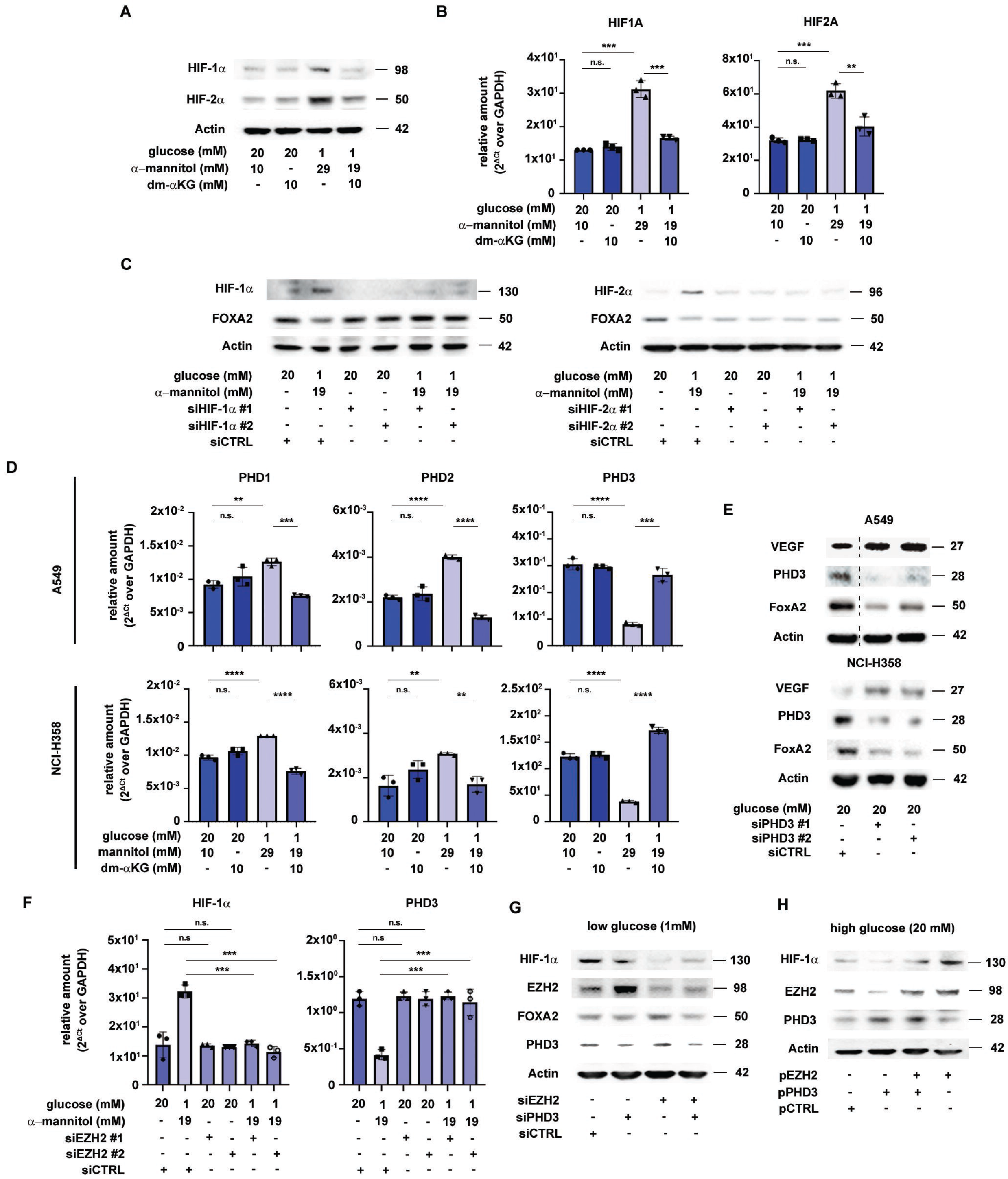

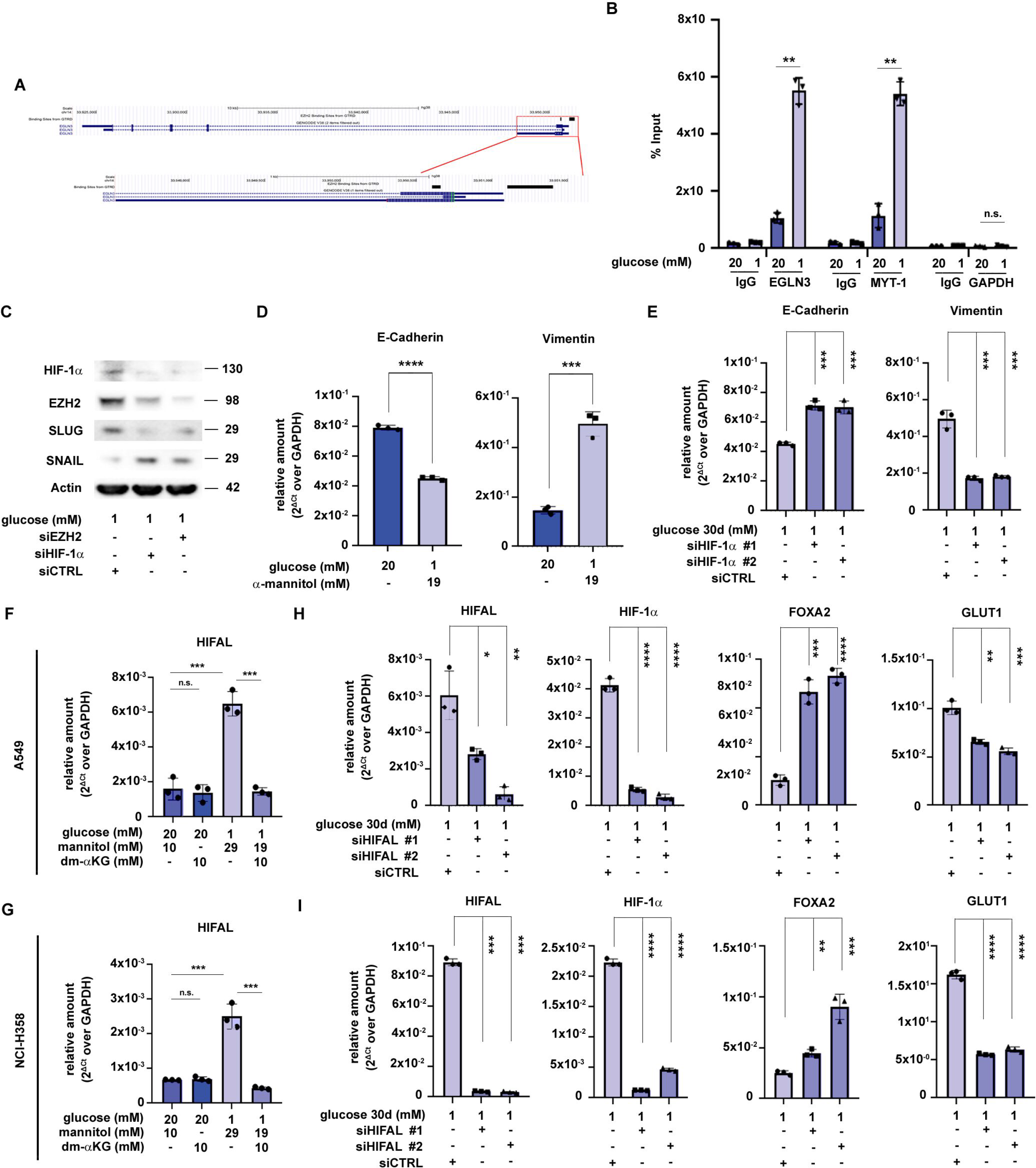

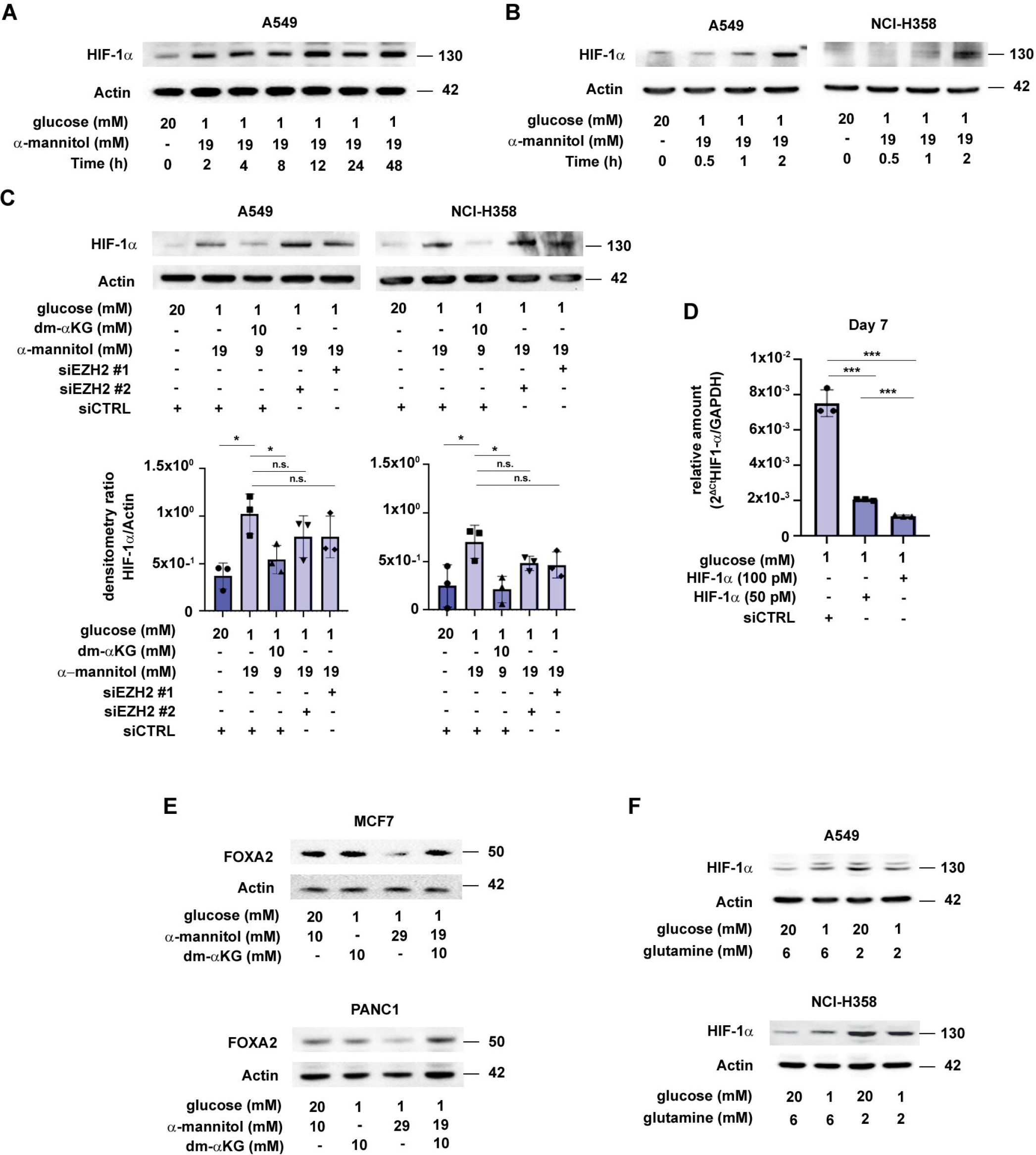

Reagents and Tools Table.

Reagents and Tools Table.
| Reagent/Resource | Reference or Source | Identifier or Catalog Number |
| --- | --- | --- |
| **Experimental models: Cell lines** |  |  |
| A549 | ATCC | CCL-185 |
| NCI-H358 | ATCC | CRL-5807 |
| Mouse: 2953A | Generated by Scafoglio Lab | N/A |
| Human: PDO | Generated by Scafoglio Lab | N/A |
| **Experimental models: Organisms/strains** |  |  |
| Gt(ROSA)26Sortm1(Luc)Kael/J | The Jackson Laboratory | JAX034320 |
| Wild type FVB/NJ | The Jackson Laboratory | 001800 |
| **Recombinant DNA** |  |  |
| pAd5CMVCreMT1pA | University of Iowa – Viral Vector Core | Cat#G0166 |
| pCMVHA hEZH2 | Addgene | Cat#24230 |
| pLp6.3-Egln3 | Addgene | Cat#79115 |
| **Antibodies (Western Blot)** |  |  |
| Rabbit mAb anti-EZH2 (1:1000) | Cell Signaling Technology | Cat#5246 |
| Rabbit mAb anti-FoxA2/HNF3β (1:2000) | Cell Signaling Technology | Cat#8186 |
| Rabbit mAb anti-Thyroid Transcription Factor 1 (TTF-1) (1:2000) | Cell Signaling Technology | Cat#12373 |
| Rabbit mAb anti-Tri-Methyl-Histone H3 (Lys4) (1:1000) | Cell Signaling Technology | Cat#9751 |
| Rabbit mAb anti-Tri-Methyl-Histone H3 (Lys9) (1:1000) | Cell Signaling Technology | Cat#9649 |
| Rabbit mAb anti-Tri-Methyl-Histone H3 (Lys27) (1:1000) | Cell Signaling Technology | Cat#9733 |
| Rabbit mAb anti-Snail (1:500) | Cell Signaling Technology | Cat#3879 |
| Rabbit mAb anti-Slug (1:500) | Cell Signaling Technology | Cat#9585 |
| Rabbit mAb anti-ZEB1 (1:1000) | Cell Signaling Technology | Cat#3396 |
| Rabbit mAb anti-Actin (1:10000) | Cell Signaling Technology | Cat#3700 |
| Rabbit pAb anti-Histone H3 (1:5000) | Cell Signaling Technology | Cat#9715 |
| Rabbit pAb anti-JMJD3 (1:1000) | Cell Signaling Technology | Cat#3457 |
| Rabbit mAb anti-UTX (1:500) | Cell Signaling Technology | Cat#33510 |
| Rabbit pAb anti-EGLN3/PHD3 (1:500) | Novus Biologicals | Cat#NB100-139 |
| Rabbit pAb anti-HIF-1α (1:500) | GeneTex | Cat#GTX127309 |
| Rabbit pAb anti-HIF-2α (1:1000) | GeneTex | Cat#GTX103707 |
| Rabbit pAb anti-VEGF (1:1000) | GeneTex | Cat#GTX102643 |
| **Antibodies (Immunohistochemistry)** |  |  |
| Rabbit mAb anti-FoxA2/HNF3β (1:400) | Cell Signaling Technology | Cat#8186 |
| Rabbit mAb anti-Thyroid Transcription Factor 1 (TTF-1) (1:1000) | Cell Signaling Technology | Cat#9751 |
| Rabbit mAb E-Cadherin (1:600) | Cell Signaling Technology | Cat#3195 |
| **Antibodies (Immunofluorescence)** |  |  |
| Mouse IgM anti-HT2-280 (1:100) | Terrace Biotech | Cat#TB-27AHT2-280 |
| Pan-CK (1:200) | Dako | Cat#GA05361-2 |
| **Antibodies (Chromatin Immunoprecipitation)** |  |  |
| Rabbit mAb anti- Histone H3 (tri methyl K4) antibody - ChIP Grade | Abcam | Cat# ab8580 |
| Rabbit mAb anti- Histone H3 (tri methyl K27) antibody - ChIP Grade | Abcam | Cat#ab195477 |
| Rabbit mAb anti-EZH2 | Cell Signaling Technology | Cat#5246 |
| **Oligonucleotides and other sequence-based reagents** |  |  |
| siRNA target sequences | This study | Table EV10 |
| RT-PCR primers | This study | Table EV11 |
| Primer sequences for ChIP RT-PCR | This study | Table EV12 |
| **Chemicals, Enzymes and other reagents** |  |  |
| Empagliflozin (BI 10773) | Selleck Chemicals | Cat#S8022;<br>CAS: 864070-44-0 |
| Tazemetostat (EPZ-6438) | MedChem Express | Cat#HY-13803<br>CAS: 1403254-99-8 |
| **Other** |  |  |
| TruSeq ChIP Library Preparation Kit | Illumina | IP-202-1012 |
| Formaldehyde | Sigma | 252549 |
| Glycine | Fisher Chemicals | G48-212 |
| PureLink RNase A | Thermo Fisher Scientific | Cat#12091021 |
| Proteinase K | New England Biolabs | Cat#P8107S |
| Dynabeads M-280 sheep anti rabbit IgG | Thermo Fisher Scientific | Cat#11203D |
| Dynabeads Protein G | Invitrogen | Cat#10003D |
| Agencourt AMPure XP | Beckman Coulter | Cat#A63882 |
| SensiFast RT Kit | Meridian Bioscience | Cat#BIO-65053 |
| iTaq™ Universal SYBR® Green Supermix | Biorad | Cat#1725121 |
| Histone H3 methylated Lys27 ELISA (H3K27) | Active Motif | Cat#53106 |
| Total Histone H3 ELISA | Active Motif | Cat#53110 |
| PureLink™ Quick Plasmid Miniprep Kit | Thermo Fisher Scientific | Cat#K210011 |
| PureLink™ HiPure Plasmid Filter Maxiprep Kit | Thermo Fisher Scientific | Cat#K210017 |
| Histone Extraction Kit | Abcam | Cat#ab113476 |
| CD326 (EpCAM) MicroBeads, mouse | Miltenyi Biotec | Cat#130-105-958 |
| MS Columns | Miltenyi Biotec | Cat#130-042-201 |
| **Deposited data** |  |  |
| RNA-Seq raw data | ArrayExpress (EMBL-EBI) | AN: E-MTAB-11253 |
| ChIP-seq raw data | ArrayExpress (EMBL-EBI) | AN: E-MTAB-11678 |
| **Software** |  |  |
| Living Image Software | Perkin Elmer | https://www.perkinelmer.com/product/spectrum-200-living-image-v4series-1-128113 |
| Aperio ImageScope software | Leica Biosystem | https://www.leicabiosystems.com/digital-pathology/manage/aperio-imagescope/ |
| TraceFinder 4.1 | Thermo Fisher Scientific | https://www.thermofisher.com/us/en/home/industrial/mass-spectrometry/liquid-chromatography-mass-spectrometry-lc-ms/lc-ms-software/lc-ms-data-acquisition-software/tracefinder-software.html |
| Trimmomatic version 0.38 | Bolger et al., 2014 | http://www.usadellab.org/cms/?page=trimmomatic |
| STAR v2.7.6a | Dobin et al., 2013 | http://code.google.com/p/rna-star/ |
| DESeq2 | Love et al., 2014 | https://bioconductor.org/packages/release/bioc/html/DESeq2.html |
| Bowtie | Langmead et al., 2009 | http://bowtie-bio.sourceforge.net/index.shtml |
| HOMER | Heinz et al., 2010 | http://homer.ucsd.edu/homer/ |
| Bedtools | Quinlan et al., 2010 | https://bedtools.readthedocs.io/en/latest/ |
| Ciider | Gearing et al., 2019 | http://ciiider.com/ |
| Ingenuity pathway analysis (IPA) | QIAGEN | https://www.qiagen.com/us/products/discovery-and-translational-research/next-generation-sequencing/informatics-and-data/interpretation-content-databases/ingenuity-pathway-analysis/ |
| Prism 8.0 software | GraphPad | https://www.graphpad.com/scientific-software/prism/ |
| QuPath | Bankhead et al., 2017 | https://qupath.github.io/ |
| Kaplan-Meier Plotter | Gyorffy et al., 2021 | https://kmplot.com/ |
| **Other** |  |  |
| TruSeq ChIP Library Preparation Kit | Illumina | IP-202-1012 |
| Glycine | Fisher Chemicals | G48-212 |
| PureLink RNase A | Thermo Fisher Scientific | Cat#12091021 |
| Proteinase K | New England Biolabs | Cat#P8107S |
| Dynabeads M-280 sheep anti rabbit IgG | Thermo Fisher Scientific | Cat#11203D |
| Dynabeads Protein G | Invitrogen | Cat#10003D |
| Agencourt AMPure XP | Beckman Coulter | Cat#A63882 |
| iTaq™ Universal SYBR® Green Supermix | Biorad | Cat#1725121 |
| Histone H3 methylated Lys27 ELISA (H3K27) | Active Motif | Cat#53106 |
| Total Histone H3 ELISA | Active Motif | Cat#53110 |
| PureLink™ Quick Plasmid Miniprep Kit | Thermo Fisher Scientific | Cat#K210011 |
| PureLink™ HiPure Plasmid Filter Maxiprep Kit | Thermo Fisher Scientific | Cat#K210017 |
| Histone Extraction Kit | Abcam | Cat#ab113476 |
| CD326 (EpCAM) MicroBeads, mouse | Miltenyi Biotec | Cat#130-105-958 |
| MS Columns | Miltenyi Biotec | Cat#130-042-201 |
| **Deposited data** |  |  |
| RNA-Seq raw data | ArrayExpress (EMBL-EBI) | AN: E-MTAB-11253 |
| ChIP-seq raw data | ArrayExpress (EMBL-EBI) | AN: E-MTAB-11678 |

